# Characterizing microbubble-mediated permeabilization in a vessel-on-a-chip model

**DOI:** 10.1101/2024.08.28.609836

**Authors:** Bram Meijlink, Gonzalo Collado Lara, Kristina Bishard, James P. Conboy, Simone A.G. Langeveld, Gijsje H. Koenderink, Antonius F.W. van der Steen, Nico de Jong, Inés Beekers, Sebastiaan J. Trietsch, Klazina Kooiman

## Abstract

Drug transport from blood to extravascular tissue can locally be achieved by increasing the vascular permeability through ultrasound-activated microbubbles. However, the mechanism remains unknown, including whether short and long cycles of ultrasound induce the same onset rate, spatial distribution, and amount of vascular permeability increase. Accurate models are necessary for insights into the mechanism so a microvessel-on-a-chip is developed with a membrane-free extravascular space. Using these microvessels-on-a-chip, we show distinct differences between 2 MHz ultrasound treatments with 10 or 1000 cycles. The onset rate is slower for 10 than 1000 cycles, while both cycle lengths increase the permeability in spot-wise patterns without affecting cell viability. Significantly less vascular permeability increase and sonoporation are induced for 10 versus 1000 cycles at 750 kPa (i.e., highest studied peak negative acoustic pressure (PNP)). The PNP threshold for vascular permeability increases is 750 versus 550 kPa for 10 versus 1000 cycles, while this is 750 versus 220 kPa for sonoporation. Vascular permeability increases do not correlate with α_v_β_3_-targeted microbubble behavior, while sonoporation correlates with α_v_β_3_-targeted microbubble clustering. In conclusion, the further mechanistic unraveling of vascular permeability increase by ultrasound-activated microbubbles in a developed microvessel-on-a-chip model aids safe and efficient development of microbubble-mediated drug transport.

## 1. Introduction

The vessel wall is an important barrier for the successful delivery of drugs to the underlying diseased tissue. An impermeable vascular wall caused by, for example, the blood-brain barrier, or as a result of anti-angiogenic treatment for non-brain cancer, hinders extravasation of pharmacological agents and thus reduces treatment efficiency.^[1–3]^ A potential way to increase treatment efficiency and reduce side-effects is to locally enhance vascular permeability using lipid-coated gas microbubbles (MBs) of 1-10 µm in diameter in combination with ultrasound.^[4,5]^

MBs have clinically been used as a contrast agent for diagnostic ultrasound imaging for several decades.^[6,7]^ Previous research has also shown that the expansion, contraction, and collapse of MBs by ultrasound can induce multiple vascular drug delivery pathways, such as sonoporation, cell-cell contact opening, and endocytosis.^[8–10]^ Sonoporation and endocytosis result in intracellular drug uptake induced by the mechanical stresses following MB oscillations close to a cell.^[10,11]^ In addition, sonoporation increases intracellular calcium,^[9,12–14]^ which correlated with cell-cell contact opening.^[15]^ As a result of oscillating MBs, the formation of an apical-to-basal tunnel through the cell has recently been reported.^[16,17]^ Cell-cell contact opening and tunnel formation could potentially induce transcellular and paracellular transportation of molecules over the endothelial layer, resulting in local MB-mediated vascular permeability increase. Investigations into vascular permeability increases by oscillating MBs are mainly studied in animal models, where both short 10 cycle^[18]^ and longer 100^[19]^ and 10,000^[20]^ cycles have shown to increase vascular permeability. However, to the best of our knowledge there has not yet been a study in non-brain vessels which directly compared the short with long cycles in the same model, either *in vitro* or *in vivo*. For vascular permeability increases outside the brain, it therefore remains unknown whether short and long cycles of ultrasound induce the same onset rate, spatial distribution, and amount of vascular permeability increase. At the same time, the reported clinical study on increasing chemotherapy delivery by oscillating MBs for pancreatic cancer, used 4 cycles.^[21]^ The reason for this short cycle is the fixed number of cycles in clinical ultrasound machines.^[22]^ This is another reason why MB-cell-drug interactions translating to vascular permeability changes need to be investigated for both short and long cycles so maximum therapeutic outcome and widespread clinical use can be achieved.

Accurate models are necessary to provide insights into the mechanism of microbubble-mediated drug delivery, thereby enabling safety and controllability. Up to now, *in vitro* research on MB-mediated vascular drug delivery pathways has been performed almost exclusively on two-dimensional (2D) endothelial cell models, in which cells are grown in a monolayer on a glass or membrane substrate.^[23]^ Consequently, it is challenging to investigate the transportation of molecules beyond the endothelial layer in the current 2D models. Moreover, 2D models usually use static growth conditions while it has been shown that growing endothelial cells under flow affects sonoporation and calcium fluxes.^[24]^ MB-mediated transportation of molecules beyond the endothelial layer under flow conditions has been investigated optically *in vivo* using chicken embryo^[18]^ and dorsal window mice models.^[20]^ While animal models can provide more accuracy, they raise ethical concerns, have a lower throughput than *in vitro* models and the differences between animals limits the controllability of the model. These considerations emphasize the need for an advanced *in vitro* model with physiological relevant cell behavior to investigate MB-mediated vascular permeability changes.

Microvessel-on-a-chip models are advanced *in vitro* models in which three-dimensional (3D) artificial blood vessels are grown under flow.^[25]^ For this reason, they are more physiologically relevant than *in vitro* 2D monolayers. At the same time, they provide a higher throughput with fewer ethical objections than *in vivo* models. Two recent studies showed the feasibility to use vessel-on-a-chip models to gain insight in microbubble-mediated sonoporation under flow^[26]^ and the induction of cell-cell contact gaps^[27]^. However, both these models still contained a membrane, meaning vascular permeability could not be investigated. The OrganoPlate^®^ 3-lane 40 (4004400B, Mimetas, Leiden, NL) is a commercially available organ-on-a-chip model in which an advanced membrane-free 3D microvessel can be grown under flow alongside an extravascular space.^[28,29]^ Acoustic characterization of the OrganoPlate^®^ indicated that controlled MB behavior can be achieved within the chips.^[30]^ These properties make the microvessel-on-a-chip grown in the OrganoPlate^®^ a promising model to investigate vascular permeability and how this permeability is affected by MB and ultrasound treatment.

In this study, we developed a microvessel-on-a-chip model with a perfused lumen and extravascular space in the OrganoPlate^®^ 3-lane with the aim to investigate ultrasound and α_v_β_3_-targeted MBs (α_v_β_3_-tMBs) mediated changes in vascular permeability and sonoporation. At increasing acoustic peak negative pressures (PNP), we investigated 2 MHz pulses of 10-cycles as well as 1000-cycles, based on previous publications that showed that the short 10-cycle^[8,9,18]^ and long 1000-cycle pulses^[26,31,32]^ can induce sonoporation, cell-cell-contact opening and vascular permeability *in vitro* and *in vivo*. The applied pulses were repeated 10 times since pulse repetitions have been shown to enhance vascular permeability *in vivo.*^[20]^ Using real-time microscopy, α_v_β_3_-tMBs dynamics were recorded, vascular permeability was investigated for 2 h with a barrier integrity assay and sonoporation was investigated with the model drug propidium iodide. From these microscopy studies, the onset rate and spatial distribution of the vascular permeability increase were determined and correlated to MB dynamics and sonoporation. The effect of the MB and ultrasound treatment on the cell viability in the microvessel was assessed using a WST-8 colorimetric assay which measures metabolic activity.

## 2. Results and Discussion

### 2.1. Establishing the microvessel-on-a-chip model

The microfluidic OrganoPlate^®^ 3-lane 40 platform (**Figure 1**A) was used to grow a 3D microvessel-on-a-chip model (Figure 1B). To achieve a membrane-free configuration between the cellular layer and extravascular space, collagen I extracellular matrix (ECM) gel was used. Depending on the lot numbers (n = 3), the median Young’s modulus of the gel measured between 0.43-0.68 kPa (**Figure S1**). These values are within previously reported ranges for endothelial cells (0.15-10 kPa)^[33]^ and orders of magnitude lower than the reported GPa Young’s modulus for polymer membranes, plastics, glass, or human bone.^[34,35]^ The observed 1.6-fold differences between the lot numbers could be a result of variation during the isolation of the collagen by the supplier, or be due to temperature differences during the neutralization and seeding of the gel.^[36]^

**Figure 1.**
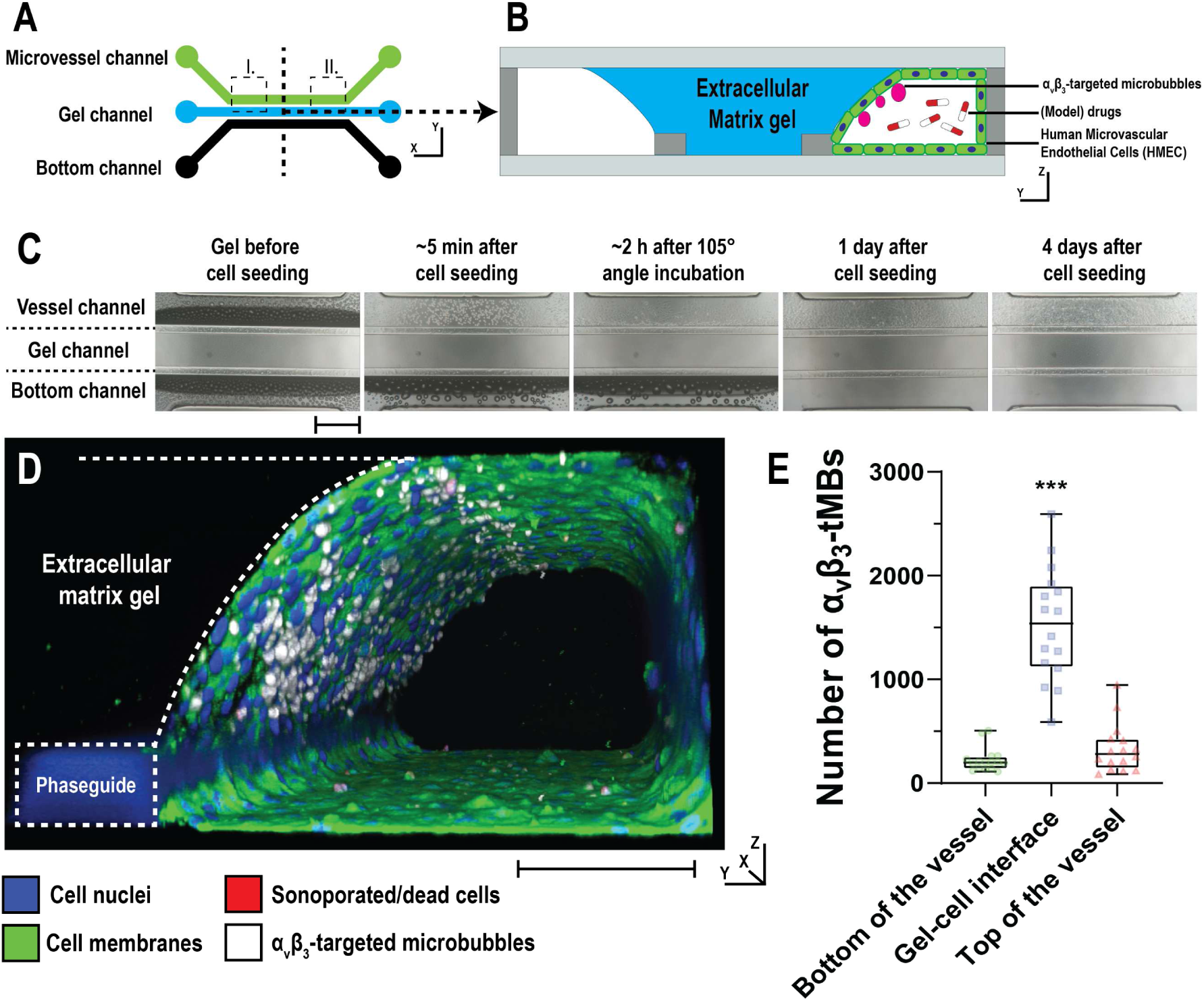
Formation of microvessel-on-chip model and live cell 3D imaging. A) Schematical representation of the three microfluidic channels from the top view with their in- and outlets. Dotted outlines I and II indicate the locations where three-dimensional confocal microscopy images were made. B) Side view of a cross-section of the microvessel-on-a-chip model as used during the experiments (not drawn to scale). The α_v_β_3_-targeted microbubbles adhered to the endothelial cell-gel interface and model drugs were added to the lumen of the vessel. C) Brightfield images of the gel seeding and vessel formation over time. Scalebar represents 500 µm. D) Three-dimensional confocal microscopy image made at location I in (A) of a live microvessel-on-chip 4 days after seeding (corresponding confocal microscopy recording animation in Video S1). Scalebar represents 100 µm for y-z direction; x-direction is 635 µm. E) Number of bound α_v_β_3_-targeted microbubbles in live microvessels-on-chip (n = 16 fields of view), including the example in D. Boxplot represents the median and the boxes indicate the 25^th^ and 75^th^ percentiles with whiskers ranging from the minimum to maximum value. Significance is indicated with *** (p<0.001). α_v_β_3_-tMBs = α_v_β_3_-targeted microbubbles.

Microvessel culture was performed based on a previous publication^[37]^. In short, microvessel growth started by dispensing the ECM gel into the middle channel (Figure 1C). Following gel loading, human dermal microvascular endothelial cells (HMEC-1) were seeded and incubated under a 105° angle to attach the cells to the gel layer. The microvessel formation within the vessel channel over a 4-day time period, in which the cells were cultured under bidirectional flow, is shown in Figure 1C. Using this method, a microphysiological 3D model was cultured which is more representative of the *in vivo* situation in comparison to glass or membrane substrates used in 2D models.^[38]^

The expression of the angiogenic biomarker α_v_β_3_ by the microvascular cells in the chip was confirmed using immunohistochemistry, albeit that non-specific staining in the ECM gel was observed for the secondary antibody in both the α_v_β_3_ and isotype control (**Figure S2**). Live cell 3D confocal microscopy imaging of a 635 µm wide section of the microvessels was performed following α_v_β_3_-tMB incubation under a 75° angle (Figure 1D and **Video S1**; α_v_β_3_-tMB in white). Quantification of 16 live cell 3D images in eight different microvessels revealed that a median of 1.5·10^3^ (interquartile range: 1.1·10^3^-1.9·10^3^) α_v_β_3_-tMBs bound to the gel-cell interface layer (Figure 1E). This was a significant 5.5-fold and 7.7-fold higher in comparison to the bound α_v_β_3_-tMBs on the top and bottom of the microvessel respectively. The α_v_β_3_-tMB thus predominantly bound to the cellular layer of interest, namely the cells located on the membrane-free gel-cell interface layer which has an extravascular space. This finding indicates that placing the OrganoPlate^®^ under a 75° angle during α_v_β_3_-tMB administration was effective in selectively targeting the α_v_β_3_-tMB by buoyancy to the cells at the gel-cell interface layer. The largest spread in the number of bound α_v_β_3_-tMB was found at the gel-cell interface, namely a 4.4-fold difference between the lowest and highest value. Further assessment showed significantly higher α_v_β_3_-tMB binding to the left side of the microvessel (location I in Figure 1A) compared to the right side (location II in Figure 1A) (**Figure S3**). This difference can be a result of the microfluidic flow direction during α_v_β_3_-tMB incubation, which was from the left to the right.

### 2.2 Barrier integrity investigation in the microvessel-on-a-chip model

To investigate vascular permeability, the leakage of a 150 kDa fluorescent FITC-dextran model drug over time was studied in a cell-free, non-treated (sham), α_v_β_3_-tMB-only, ultrasound only (2 MHz, 750 kPa; 10×1000 cycles) and ultrasound (2 MHz, 750 kPa; 10×10 or 10×1000 cycles) plus α_v_β_3_-tMB treated microvessels (typical fluorescent image examples in **Figure 2**A). In the cell-free chips, the FITC-dextran already leaked into the bottom vessel before the first image was made, indicating the absence of a barrier. For the microvessels, the FITC-dextran gradually leaked similarly into the gel and bottom channel in the initial state, i.e., before treatment, as also quantified in Figure 2B. In the initial state, the leakage in these microvessels was below 50%. To ensure similarity in barrier function between the studied microvessels, it is common to exclude microvessels that have an insufficient barrier function^[28]^ due to, for example, differences in gel seeding or cell growth.^[29,39]^ We therefore excluded microvessels when the leakage exceeded 50% within the initial state. This resulted in the exclusion of 34% of all grown microvessels. Of these excluded microvessels, 71% were part of the section of the OrganoPlate^®^ in which the BI assay was performed first. Another Organoplate^®^ study using transepithelial electrical resistance as readout for permeability, suggested that factors like flow and liquid level change can induce permeability changes at the start of the assay which can recover over time^[40]^. The higher initial leakage observed in the first section of the OrganoPlates^®^ in our study, is therefore likely caused by moving the plate from the incubator to the experimental setup as this changes the flow and liquid levels. At the same time, the observed increase in leakage in the initial state was far less pronounced in the second section of the OrganoPlat^e®^, indicating recovery. For future studies, it may therefore be beneficial to wait with starting the BI assay until some time after the plate has been moved.

**Figure 2.**
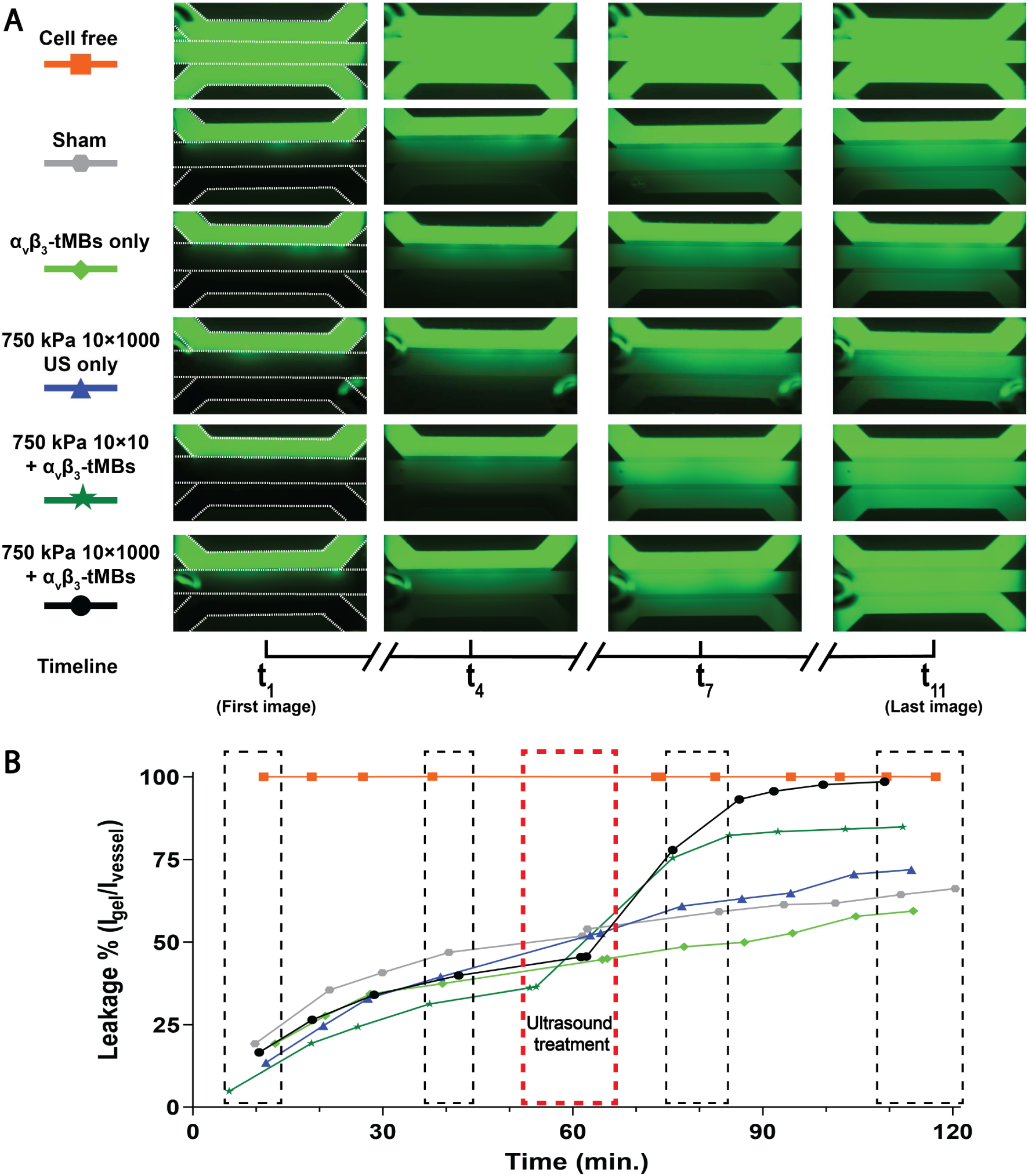
Barrier integrity assay for six different treatment conditions. A) Four selected representative images from the fluorescent microscopy sequence of images visualizing the 150 kDa FITC-dextran model drug leakage over a 2-h period. The three channels in the chip (from top to bottom: microvessel channel, gel channel, bottom channel) are outlined with white-dotted lines in the most left images. B) Leakage % (i.e., Intensity_gel_/ Intensity_vessel_) over time quantified from the sequence of fluorescent microscopy images. The black-dotted rectangles indicate the time in which the four images shown in A were recorded. The red dotted rectangle indicates the ultrasound treatment (2 MHz; 750 kPa peak negative pressure and 10×10 or 10×1000 cycles) for the conditions US only and US plus α_v_β_3_-targeted microbubbles (α_v_β_3_-tMBs).

After treatment, the leakage in the sham, α_v_β_3_-tMB-only, and ultrasound only conditions was unaffected, resulting in a final leakage percentage (median with interquartile range and n-number between brackets) of ∼73% (62%-83%; n=36) at the end of the BI assay. By contrast, a clear increase in leakage was observed for the ultrasound plus α_v_β_3_-tMB conditions after treatment. At the end of the BI assay, the leakage was ∼89% (85%-92%; n=8) for the 750 kPa 10×10 cycles plus α_v_β_3_-tMB condition, while this was ∼98% (93%-100%; n=9) for the 750 kPa 10×1000 cycles plus α_v_β_3_-tMB condition. Following 0.1% Triton-X addition, the leakage percentage of all microvessels was 100% (100%-100%; n=122), which indicates that the microvessels having a leakage of ∼89% at the end of a BI assay still had a barrier.

### 2.3 The effect of acoustic pressure and pulse length on vascular apparent permeability

The vascular apparent permeability (P_app_) was calculated before and after treatment from the obtained leakage patterns in the BI assay. Four control (sham; α_v_β_3_-tMBs only; 750 kPa 10×10 cycles ultrasound only; 750 kPa 10×1000 cycles ultrasound only) and ten different ultrasound plus α_v_β_3_-tMBs treatment conditions (5 different *in situ* acoustic pressures ranging from 90– 750 kPa at 10×10 or 10×1000 cycles) were investigated in a total of 122 microvessels, in six different OrganoPlates^®^. In the initial state (i.e., before treatment), only one significant difference in P_app_ was observed, namely between the sham and the 750 kPa 10×1000 cycles ultrasound only condition (**Figure S**4, all p-values in **Table S**1). This significant difference likely occurred due to differences in vessel-growth during culture.^[39]^ To address this difference between the control conditions in the initial state, the post-treatment increases in vascular permeability were considered significant when different compared to all four control conditions. After ultrasound plus α_v_β_3_-tMB treatment, a significant increase in P_app_ was observed for the highest applied acoustic pressure of 750 kPa at both 10×10 cycles and 10×1000 cycles and the second highest applied acoustic pressure of 550 kPa at 10×1000 cycles in comparison to the control treatments (**Figure 3**; all p-values in **Table S**2). Specifically, in comparison to the ultrasound only control condition of 750 kPa and 10×1000 cycles, the P_app_ for the ultrasound plus α_v_β_3_-tMBs treated microvessels was ∼2.4-fold higher for 750 kPa and 10×10 cycles while this was ∼3.5-fold higher for 550 kPa and 10×1000 cycles and ∼5-fold higher for 750 kPa and 10×1000 cycles. In addition, the P_app_ for microvessels treated with α_v_β_3_-tMBs at 750 kPa and 10×10 cycles was a significantly 2.5-fold lower than at 750 kPa and 10×1000 cycles. This shows that higher ultrasound pressures increasingly enhanced the vascular permeability upon α_v_β_3_-tMB-mediated treatment, which is in line with what others have observed *in vivo* using chicken embryo^[18]^ and mouse cranial^[19]^ and tumor window^[20,41]^ models. A previous vessel-on-a-chip study showed significantly more cell-cell junction opening was induced with ultrasound pulses with an MI of 0.72 compared to an MI of 0.4 (600×500 cycles), which could explain why leakage in our model was only observed with the higher ultrasound pressures.^[27]^

**Figure 3.**
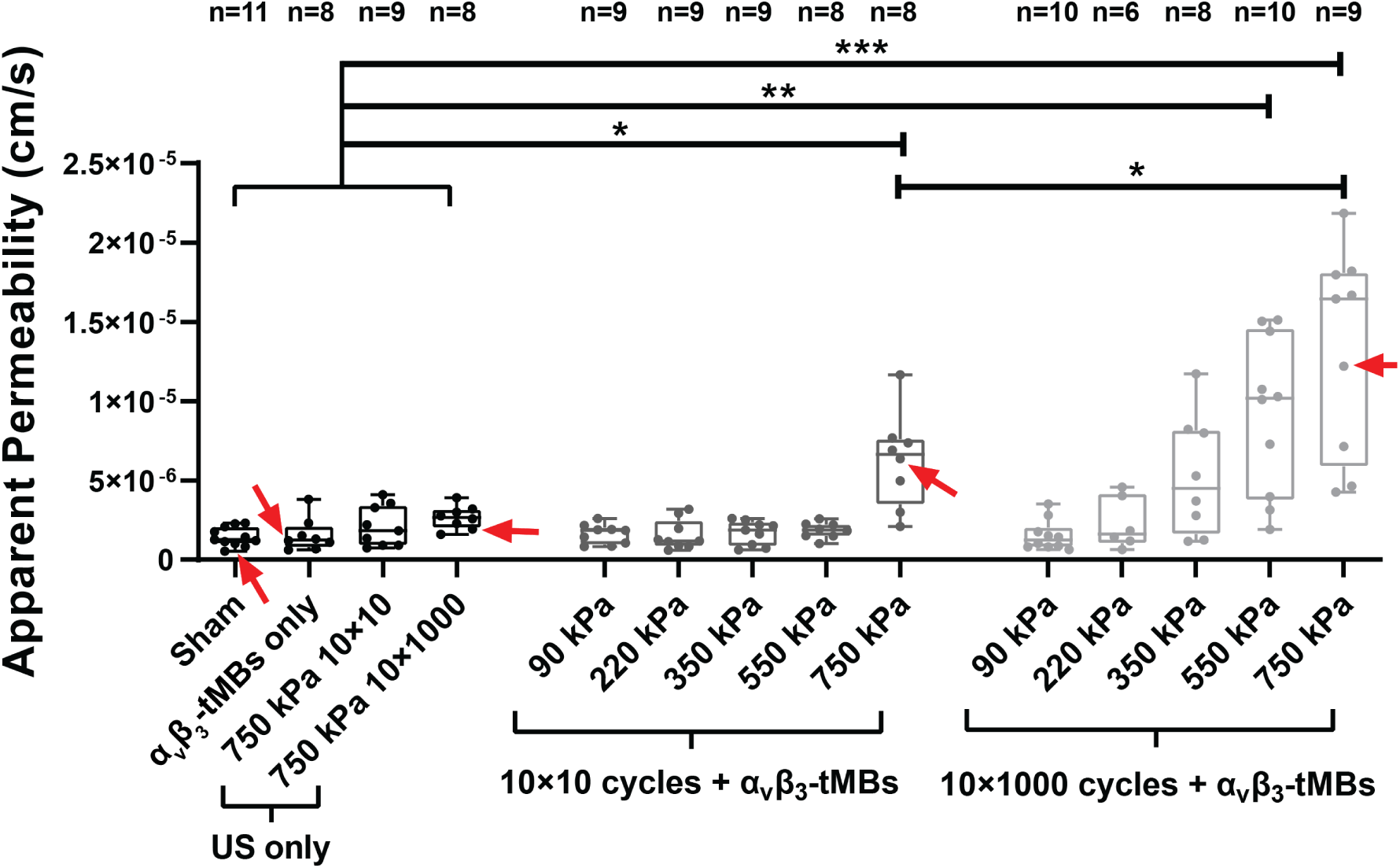
Apparent permeability after treatment for four control (on left) and ten treatment conditions. Boxplot represents the median with the boxes indicating the 25^th^ and 75^th^ percentiles with whiskers ranging from the minimum to maximum value. Significance between the four control and ten treatment conditions is indicated with *p<0.05, **p<0.01, and ***p<0.001. See Table S2 for all statistical comparisons. Datapoints that correspond to the example vessels shown in Figure 2 are indicated with red arrows. US = ultrasound; α_v_β_3_-tMBs = α_v_β_3_-targeted microbubbles.

In addition, our findings that bursts of just 10 cycles can already increase the vascular permeability are in agreement with a previous *in vivo* chicken embryo study^[18]^. The minimum pressure to induce permeability in our study using 2 MHz ultrasound was 750 kPa for 10×10 cycles (i.e., mechanical index (MI) = 0.53) plus α_v_β_3_-tMBs and 550 kPa for 10×1000 cycles (i.e., MI = 0.39) plus α_v_β_3_-tMBs. These MI values are in agreement with the minimum MI to induce permeability in mouse cranial window^[19,42]^ (i.e., MI = 0.37 with 600×100 cycles and MI = 0.55 with 120×12000 cycles) studies and a tumor window^[20]^ study (i.e., MI = 0.4 with 150×1000 cycles). On the other hand, the MI of 0.53 needed to induce permeability for the shorter 10×10 cycle treatment plus α_v_β_3_-tMBs in our study was considerably lower than the previously reported needed MI of 1.3 for the 2500×10 cycle plus MB treatment in a chicken embryo study^[18]^. A plausible explanation for this difference is the smaller vessels with diameters of 12-21 µm that were studied in the chicken embryo in comparison to 200-300 µm vessels in the OrganoPlate^®^ 3-lane, as other studies have reported that the MB’s oscillation amplitude is 2 – 3 fold less in 25 than 160 – 200 µm diameter capillary tubes^[43,44]^ due to the closer proximity of the opposite wall to the MBs.^[43]^

### 2.4 Onset rate and spatial distribution of the vascular permeability

For the three ultrasound plus α_v_β_3_-tMB treatment conditions that showed a significant increase in P_app_ in comparison to the four control conditions (see Figure 3), the onset rate of the leakage was further investigated during the first 5 min after treatment using time-lapse imaging as presented in **Figure 4**. For the sham and 750 kPa 10×1000 cycles ultrasound only control treatments, no increases in leakage were observed in the microvessels during the first 5 min after treatment (Figure 4A and zoom-in in Figure 4B), which is in line with the findings presented in Figure 2. We hardly observed any increase in leakage during the first 5 min after treatment for the 750 kPa 10×10 cycles plus α_v_β_3_-tMB treatment, while the mean leakage percentage did reach 71% at the end of the BI assay (Figure 4A and zoom-in in Figure 4B). By contrast, for the 550 and 750 kPa 10×1000 cycles ultrasound plus α_v_β_3_-tMBs treatment conditions, a clear increase in leakage was observed during the first 5 min after treatment (Figure 4A and zoom-in in Figure 4B). At the end of the BI assay, the mean leakage percentage reached 96% for the 550 kPa and 97% for the 750 kPa 10×1000 cycles plus α_v_β_3_-tMB treated vessels. The increase within the first 5 min upon treatment was lower and the onset rate 3-fold slower for the 550 kPa than the 750 kPa plus α_v_β_3_-tMB condition. This is in line with previous *in vivo* studies that reported significantly faster extravasation rates for higher acoustic pressures in a chicken embryo model (700 versus 1300 kPa; 10 cycles at 1 MHz)^[18]^ and faster extravasation onset times for higher acoustic pressures in a mouse tumor window model (200 versus 800 kPa; 10,000 cycles at 1 MHz).^[20]^ In our study, the calculated P_app_ showed no significant differences between the treatment conditions in the initial state, while the P_app_ during the first 5 min after treatment (Figure 4C) showed a similar trend as the P_app_ for 1.5 h after treatment (Figure 3), although not significant.

**Figure 4.**
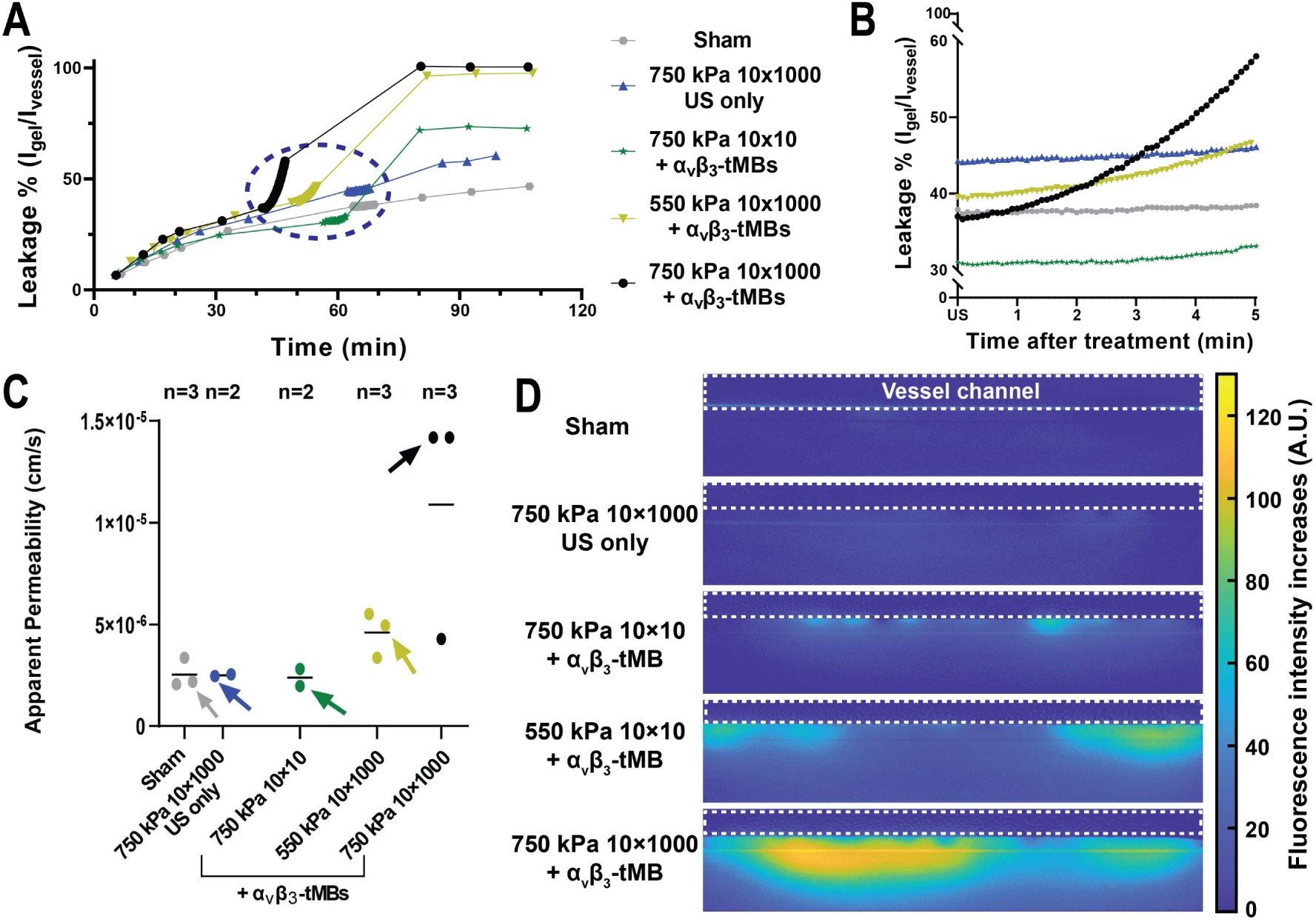
Onset rate and spatial distribution of the vascular permeability increase. A) Leakage % (Intensity_gel_/ Intensity_vessel_) over time with the integrated 5-min time-lapse imaging in the blue-dotted circle. B) Zoom-in of the blue-dotted circle of Figure 4A showing the leakage % during the 5-min time-lapse imaging directly after treatment. C) Apparent permeability (P_app_) during the first 5 min after treatment. Datapoints represent individual vessels, with arrows indicating the datapoints corresponding to the examples shown in Figure 4A, B & D, and lines indicate the mean per condition. No statistically significant differences were observed. D) Visualization of the spatial distribution of the leakage 5 min after treatment. US = ultrasound; α_v_β_3_-tMBs = α_v_β_3_-targeted microbubbles; A.U. = arbitrary unit.

The 5-min time-lapse recordings showed that the vascular permeability increase started in a spot-wise pattern for all the ultrasound plus α_v_β_3_-tMB conditions (Figure 4D and **Video S2**), including the 750 kPa 10×10 cycle plus α_v_β_3_-tMB condition that hardly had an increase in leakage percentage and P_app_. The spot-wise pattern is in line with previous chicken embryo^[18]^ and mouse model^[20]^ studies where the ultrasound plus MB-induced leakage also originated from single spots. In our study, we observed that the initial leakage spots were unevenly distributed over the microvessel and varied in number per microvessel, which could be a result of the previously observed differences in microbubble concentrations in Figure 1E.

### 2.5 The effect of the microbubble-mediated treatment on sonoporation

Before treatment with 750 kPa 10×1000 cycle ultrasound plus α_v_β_3_-tMBs, hardly any propidium iodide (PI) was observed in the 3D confocal microscopy image of part of the microvessel (**Figure 5**A), suggesting the number of dead cells was negligible. One and a half h after treatment, an increase in PI signal was mainly observed in the cells located at the gel-cell interface layer (Figure 5A). As PI stains dead^[45]^ and alive sonoporated^[9]^ cells by entering the cells and becoming fluorescent upon intercalation with RNA and DNA,^[46]^ the distinguishment between dead or sonoporated cells cannot be made 1.5 h after treatment. However, this difference can be made up to 120 s after treatment as membrane pores remaining open for more than 120 s do not close,^[9,31,47]^ consequently resulting in cell death.^[31]^ In the initial state, the only significant difference in PI signal was between the sham and lower 550 kPa 10×1000 cycles plus α_v_β_3_-tMBs treatment (**Figure S5**, all p-values in **Table S3**). Figure 5B shows the change in PI signal within 45 s after treatment. As for the P_app_, post-treatment increases were considered significant when different compared to all four control conditions. For the 10×10 cycles plus α_v_β_3_-tMB conditions, the change in PI signal was only significantly higher for the 750 kPa pressure (2.9% increase) (all p-values in **Table S4**). For the 10×1000 cycles plus α_v_β_3_-tMB conditions, a significant change in PI signal was observed for the four highest pressures: 2.5% increase at 220 kPa, 7.3% increase at 350 kPa, 24.5% increase at 550 kPa and 28.5% increase at 750 kPa. In addition, the 550 and 750 kPa 10×1000 cycles plus α_v_β_3_-tMB conditions also showed a significantly higher change in PI in comparison to all other ultrasound plus α_v_β_3_-tMB treatment conditions. Our findings are in line with two other *in vitro* studies,^[31,48]^ which showed that the higher the acoustic pressure was, the more cells were sonoporated. These studies used 1 MHz ultrasound with pressures ranging from 100-500 kPa at 2000 cycles^[48]^ or 140-500 kPa at 500-50000 cycles.^[31]^ The latter study also reported an increase in sonoporation for longer cycle treatments,^[31]^ which is also in line with our findings.

**Figure 5.**
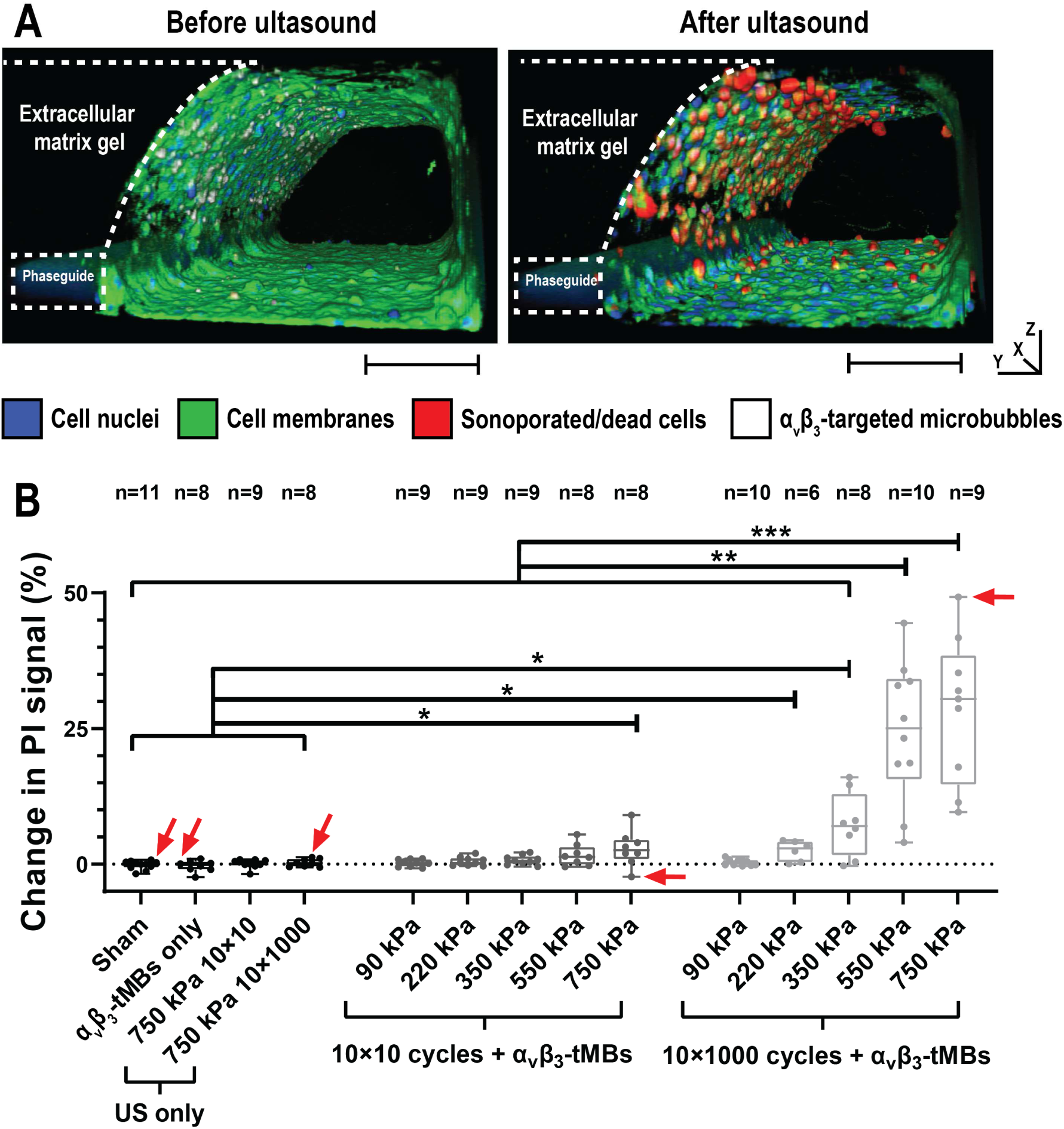
Sonoporation of cells in microvessel-on-chip after treatment. A) Three-dimensional confocal microscopy image of part of a microvessel before (left) and 1.5 h after 750 kPa 10×1000 cycles ultrasound plus α_v_β_3_-tMBs treatment (right). Scalebar represents 100 µm for y-z direction; x-direction is 635 µm. B) Percentage change in PI signal within 45 s after treatment. Boxplot represent the median and the boxes indicate the 25^th^ and 75^th^ percentiles with whiskers ranging from the minimum to maximum value. Statistical significance is indicated with *p<0.05, **p<0.01, ***p<0.001. See Table S4 for all statistical comparisons. Datapoints which correspond to the example vessels shown in Figure 2 are indicated with red arrows. PI=propidium iodide; US = ultrasound; α_v_β_3_-tMBs = α_v_β_3_-targeted microbubbles.

### 2.6 The effect of the ultrasound pulse length on microbubble behavior

The optical bright field recordings showed that the bound α_v_β_3_-tMB displaced during the 10 ultrasound pulses. For the 750 kPa 10×10 cycle condition, shown in the left panels of **Figure 6**A-C and **Video S3**, α_v_β_3_-tMBs displaced at the far right of the vessel channel near the gel-cell interface layer, while displacement occurred throughout the vessel channel for the 750 kPa 10×1000 cycle condition (right panels of Figure 6A-C and **Video S4**). The displacement of tMBs bound to a surface by the application of ultrasound has been observed before,^[31,49,50]^ and is caused by secondary Bjerknes forces, i.e., the mutual interaction between oscillating MBs. Following the displacement of the α_v_β_3_-tMBs, the formation of MB clusters was observed. In the 10×10 cycles condition, a total of four clusters formed divided over the 46 microvessels, whereas a total of 207 clusters formed divided over the 48 microvessels in the 10×1000 cycles condition (Figure 6D, **Table S5**). This difference suggests that cluster formation predominantly occurred after longer cycle insonification, which is in line with what others have observed.^[49–51]^ In the OrganoPlate^®^ 3-lane, cluster formation occurred more often at the left and right side of the microvessel close to the ends of the PhaseGuides^®^ (Figure 6D). However, no pattern in the distance between clusters was found (**Figure S6**), thereby indicating that standing waves^[52]^ were not the main mechanism leading to tMB clustering. Instead, it is more plausible that the tMB clusters were induced by secondary Bjerknes forces as these forces have previously been shown to induce MB clusters upon ultrasound insonification in a 200 µm capillary.^[53]^ In addition, others have reported that the number of formed MB clusters increased upon longer ultrasound insonification time.^[54]^

**Figure 6.**
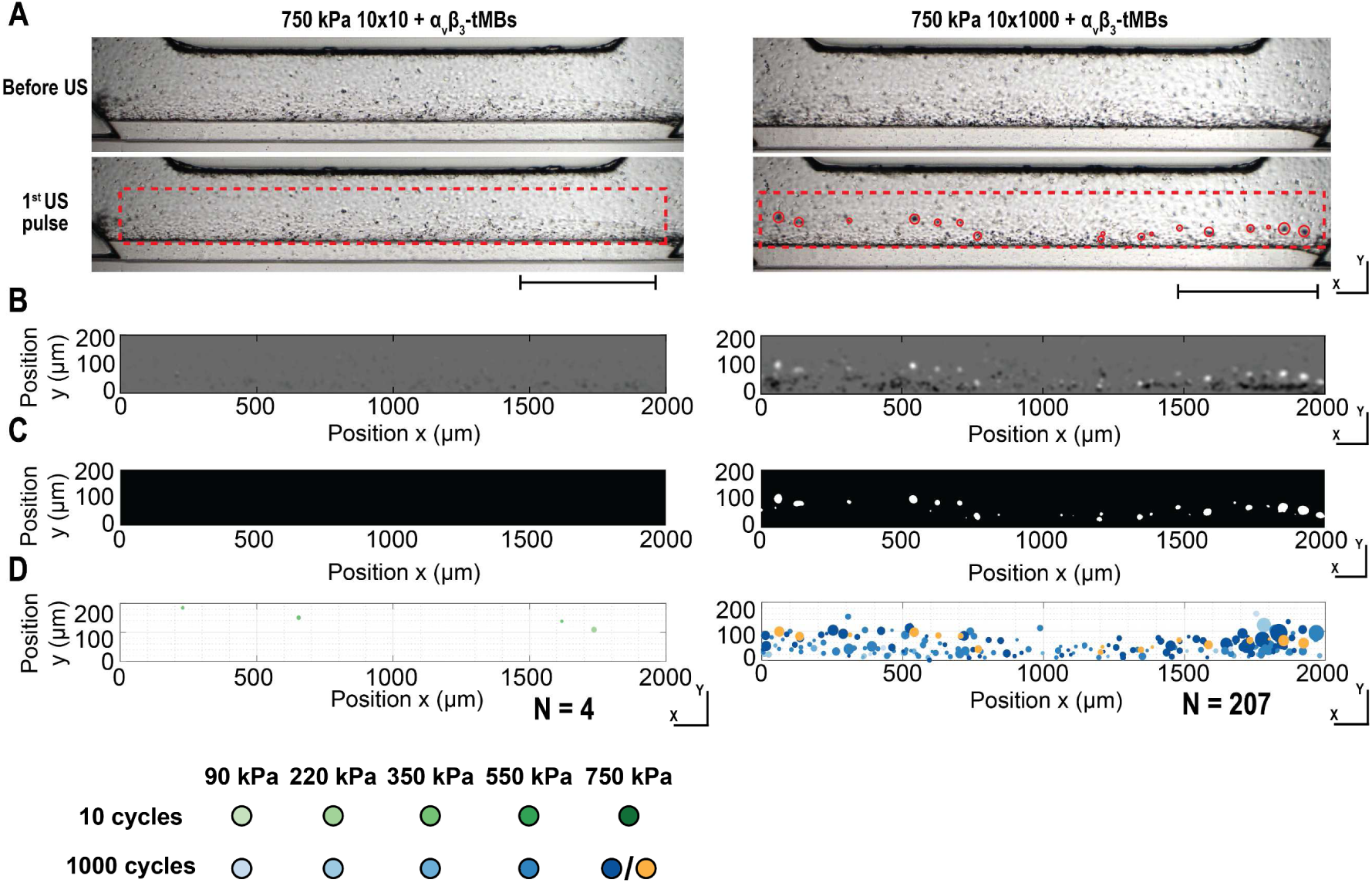
α_v_β_3_-Targeted microbubble cluster formation during ultrasound treatment. A) Example images of the microvessels from Figure 4 before ultrasound (US) (top row) and after the first US pulse (bottom row). Red-dotted rectangles indicate the ROI for the cluster analysis and red circles highlight the formed α_v_β_3_-tMB clusters. Scalebar represents 500 µm. B) Difference in signal after the first ultrasound pulse with black indicating signal increase (i.e., more light goes through) as a result of tMB displacement and white indicating signal decrease (i.e., less light gets through) as a result of tMB cluster formation. C) Mask based on the signal decrease from Figure 6B. D) Schematic representation of all tMB cluster locations and sizes obtained from 48 (left) and 46 (right) microvessels. Circle locations, sizes and colors correspond with the cluster locations, cluster sizes and treatment conditions. The orange points in the 1000 cycles graph correspond with the tMB clusters in Figure 6A. (A-D) Microvessels treated with 750 kPa plus α_v_β_3_-tMB at 10×10 cycles (left) and 10×1000 cycles (right).

### 2.7 Relation between vascular permeability, sonoporation, and *α*_v_*β*_3_-tMB clustering

The relation between α_v_β_3_-tMB cluster formation and α_v_β_3_-tMB displacement, vascular permeability increase (i.e. leakage increase), and sonoporation (i.e. PI increase), was evaluated by segmenting the treated microvessels in 200 µm segments. Only the 550 and 750 kPa 10×1000 cycles plus α_v_β_3_-tMB conditions were selected for the analysis because these conditions showed a significant increase in P_app_ (Figure 3) and contained 86% of all formed α_v_β_3_-tMB clusters (Table S5). Segments with higher α_v_β_3_-tMB displacement after the first ultrasound pulse showed significantly more cluster formation (**Figure 7**A). As shown in Figure 7B, no relation was found between the number of α_v_β_3_-tMB clusters and vascular permeability increase. This finding is in line with others who reported that MB behavior could not predict cell-cell contact opening^[9]^. By contrast, more α_v_β_3_-tMB cluster formation resulted in significantly more sonoporation within the same vessel segment after the first ultrasound pulse (Figure 7C). Other *in vivo* work also showed that MB clusters sonoporated more cells than single MBs and the sonoporation rate increased with bigger MB clusters,^[55]^ thereby validating the robustness of our model. For the applied consecutive ultrasound pulses, the number of α_v_β_3_-tMB clusters decreased suggesting the clusters moved, merged and/or dissolved (Figure S6). The consecutive pulses showed less significant relations between α_v_β_3_-tMB clusters, α_v_β_3_-tMB displacement, and sonoporation (**Figure S7**). A possible explanation for the observed differences between the first and later applied cycles of ultrasound is that single microbubbles have a higher oscillation amplitude at the applied ultrasound frequency^[51]^ than clusters of MBs, which behave as one big MB having a lower resonance frequency.^[54]^ To verify this possible explanation, ultra-high-speed imaging can be used as this technique has been shown to determine oscillation amplitudes of both single MBs^[8,9]^ and MB clusters^[56]^. A limitation of our study is that ultra-high-speed imaging was not feasible to visualize the MB oscillations due to technical challenges as a combination of an objective with a long working distance, high resolution, and high magnification is needed. Instead, passive cavitation detection^[57]^ could be used to gain insight into the response of the MB population within the microvessel-on-a-chip model.

**Figure 7.**
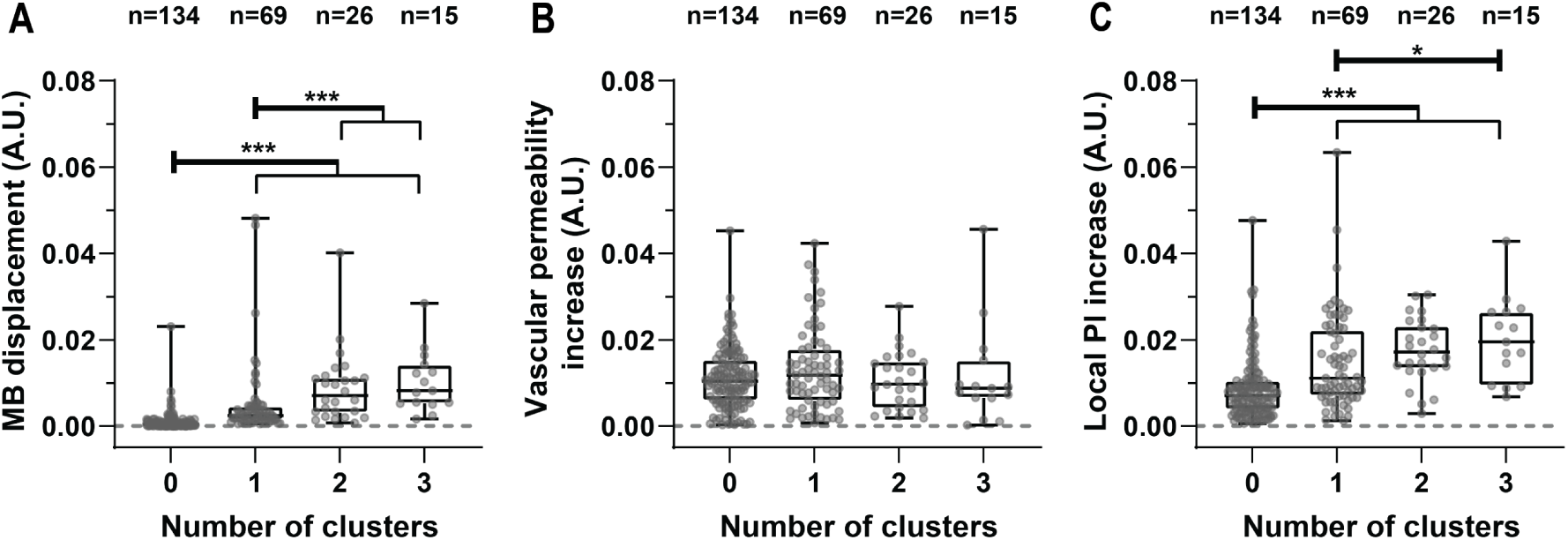
Local relations with the number of formed α_v_β_3_-tMB clusters per 200 µm segments in the microvessels after the first cycle of ultrasound. A) Local relation between formed α_v_β_3_-tMB clusters and α_v_β_3_-tMB displacement. B) Local relation between formed α_v_β_3_-tMB clusters and vascular permeability increase. C) Local relation between formed α_v_β_3_-tMB clusters and propidium iodide (PI) increase (i.e., sonoporation). (A-C) Boxplots represent the median and the boxes indicate the 25^th^ to 75^th^ percentiles with whiskers ranging from the minimum to maximum values. Every data point and n-number represent individual microvessel segments of 200 µm. Statistical significance is indicated with *p<0.05 and ***p<0.001. MB = α_v_β_3_-targeted microbubble; A.U. = arbitrary unit; PI=propidium iodide.

Further investigation into the relation between vascular permeability increases and sonoporation per microvessel segment (**Figure 8**) showed no significant correlation above the control threshold (**Figure S8**) for any of the treatment conditions. The absence of this correlation is underlined by the 220 kPa and 350 kPa 10×1000 cycles plus α_v_β_3_-tMB treatments, which both resulted in no significant vascular permeability increase (Figure 3) while sonoporation was significantly increased (Figure 5B). Our findings therefore suggest that an increase in vascular permeability is induced in microvessels in which sonoporation also occurred, while sonoporation can also be induced in the absence of an increase in vascular permeability. An increase in vascular permeability can be a result of trans-endothelial tunnel formation upon sonoporation,^[16]^ which creates transcellular gaps,^[58]^ and/or cell-cell junction opening,^[8,59]^ which creates paracellular gaps. Whether the microbubble-mediated vascular permeability increase in the by us used microvessel-on-a-chip model was due to tunnel formation and/or cell-cell contact opening is currently unknown as it is technically challenging to investigate these processes in a 3D model of this size. In our study, we used cell medium as fluid in the lumen of the microvessel. For future studies, whole blood could be used instead of cell culture medium to investigate the attribution of for example red blood cells to drug delivery as red blood cells have been shown to contribute to the microbubble-mediated induced bioeffects *in vivo.*^[60]^ At the same time, the effect of the increase in viscosity for whole blood in comparison to cell culture medium needs to be investigated with regards to the MB behavior, including MB clustering, vascular permeability increases, and sonoporation.

**Figure 8.**
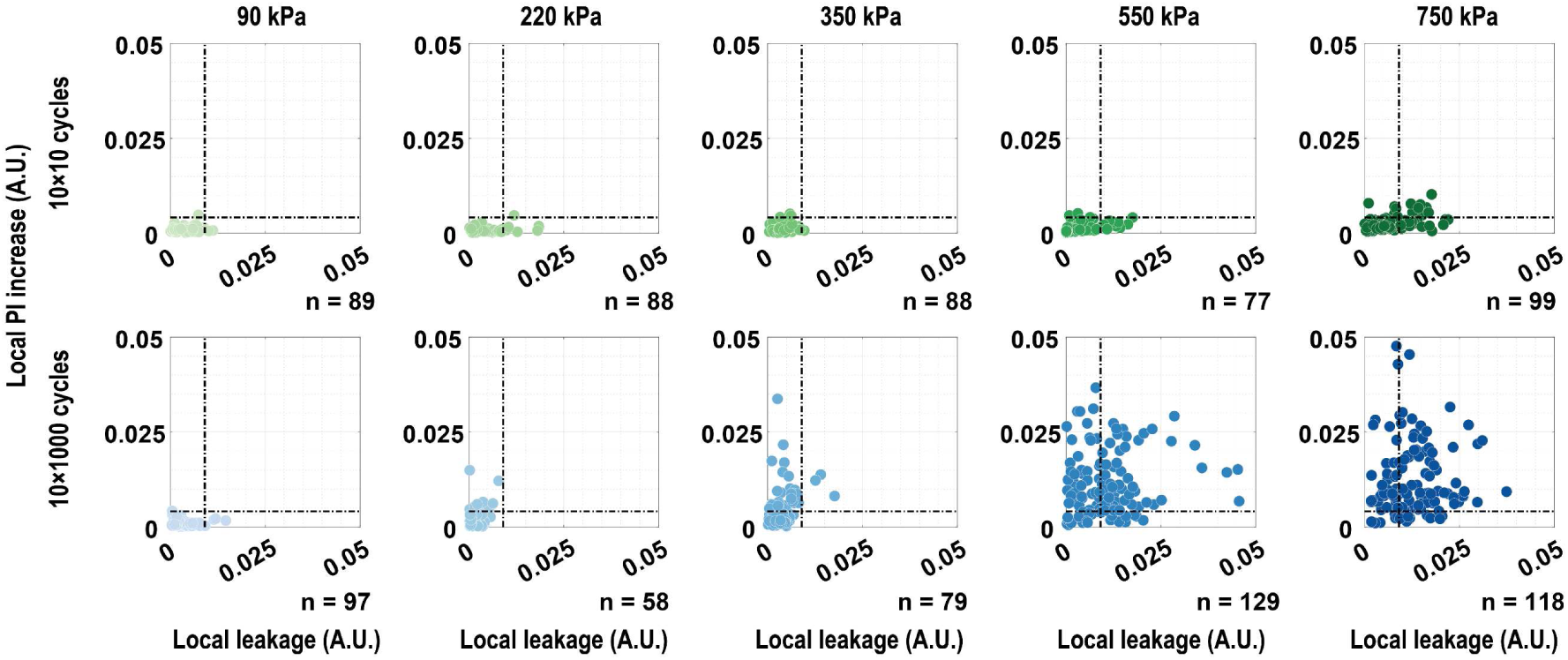
Correlation between vascular permeability increase (i.e., leakage) and sonoporation (i.e., propidium iodide uptake) in the microvessels for the ultrasound plus α_v_β_3_-tMB treatments. The top row is the 10×10 cycle treatment and bottom row the 10×1000 cycle treatment with colors corresponding to the treatment conditions as illustrated in Figure 6D. Every data point and n-number represent individual microvessel segments of 200 µm. The dotted lines indicate the thresholds obtained from the control conditions as presented in Figure S8. No statistically significant correlations were observed. PI=propidium iodide; A.U. = arbitrary unit.

### 2.8 The effect of the microbubble-mediated treatment on cell viability

The effect of the ultrasound plus α_v_β_3_-tMB treatment on the cell viability 78 min after treatment was investigated in 60 cell-containing microvessels divided over 9 different treatment conditions and 1 positive and 4 negative controls using a WST-8 colorimetric assay, as presented in **Figure 9**. The positive and negative controls had a significantly lower mean absorbance value of ∼0.3 in comparison to 0.8-0.9 for all treated cell-containing microvessels (all p-values in **Table S**6). No significant differences in cell viability were observed between all the sham, α_v_β_3_-tMB only, ultrasound only, and ultrasound plus α_v_β_3_-tMB treated microvessels, indicating that the cell viability of the entire microvessel was not affected by the treatment. Yet, ∼4% cell death has previously been reported for endothelial cells treated with ultrasound (1 MHz, 500 kPa (MI 0.5), 1×1000 cycles) plus CD31-tMB.^[31]^ The reason for the difference in cell viability between their 2D *in vitro* and our 3D *in vitro* study could be due to differences in the stiffness of the substrate on which the cells were grown, namely a polymer membrane in their study and soft collagen (Young’s modulus 0.43-0.68 kPa) in our study. This explanation is supported by a previous study that reported less ultrasound plus MB-induced cell death for cells cultured on a soft substrate (Young’s modulus 0.2 kPa) in comparison to a rigid substrate (Young’s modulus 40 kPa).^[61]^

**Figure 9.**
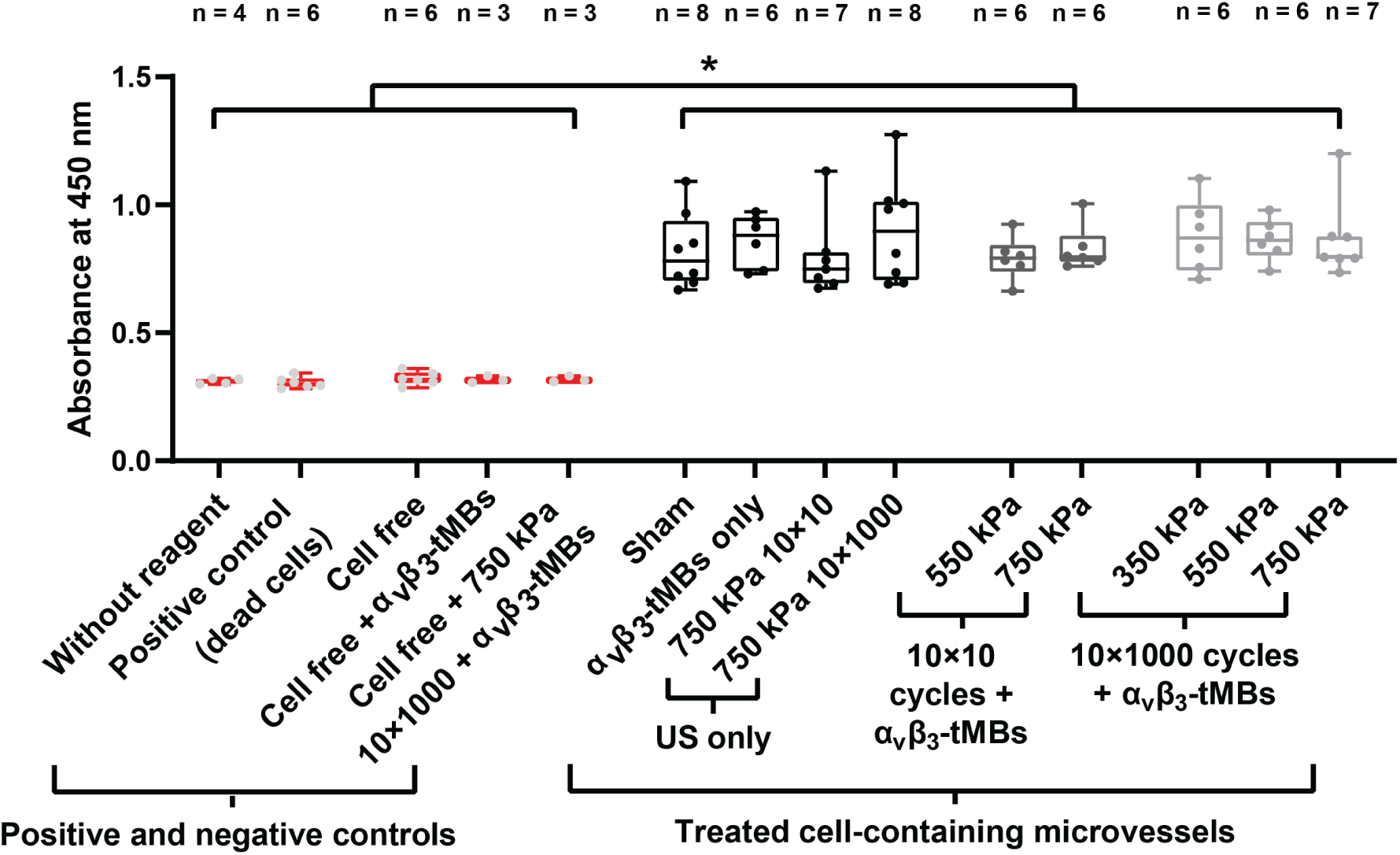
Cell viability after treatment measured with a WST-8 colorimetric assay. Boxplots represent the median and the boxes indicate the 25^th^ and 75^th^ interquartile range with whiskers ranging from the minimum to maximum value. A higher absorbance indicates more living cells. Statistical significance is indicated with *p<0.05. See Table S6 for all statistical comparisons. US = ultrasound; α_v_β_3_-tMBs = α_v_β_3_-targeted microbubbles.

## 3. Conclusions

Investigating α_v_β_3_-tMB-mediated changes in vascular permeability and sonoporation in a newly developed microvessel-on-a-chip model with a membrane-free extravascular space revealed distinct differences between ultrasound treatments with 10×10 and 10×1000 cycles. Although both the 10×10 cycles and 10×1000 cycles of 2 MHz ultrasound induced vascular permeability increases in spot-wise patterns and sonoporation without affecting cell viability, the 10×10 cycles induced significantly less vascular permeability increase and sonoporation in comparison to the 10×1000 cycles at the highest studied PNP of 750 kPa. In addition, the PNP threshold for inducing a vascular permeability increase and sonoporation was higher for 10×10 than for 10×1000 cycles, making a treatment with longer cycles of ultrasound plus α_v_β_3_-tMBs more potent for future therapeutic applications. At the same time, if α_v_β_3_-tMB-mediated drug delivery to endothelial cells via sonoporation is only needed, then lower pressures should be used as 220 and 350 kPa at 10×1000 cycles induced significant sonoporation in the absence of a vascular permeability increase. Evaluating the microbubble behavior revealed no relation with vascular permeability increases while a relation was found between formed α_v_β_3_-tMB clusters and sonoporation. This developed microvessel-on-a-chip model and the novel insights generated with it demonstrate the effect of the ultrasound pressure and cycle length on the vascular permeability increase and sonoporation outcome, which aids towards the safe and efficient implementation of MB-mediated local drug delivery.

## 4. Experimental Section

### Cell Culture

Immortalized human dermal microvascular endothelial cells (HMEC-1, CRL-3242, ATCC, Manassas, VA USA) were cultured in MCDB131 medium (10372019, Gibco, Thermo Fisher Scientific, Waltham, MA, USA). This basal medium was supplemented with 10% heat-inactivated fetal bovine serum (FBS, 16122563, Gibco), 10 mM L-Glutamine (G7513, Sigma-Aldrich, Zwijndrecht, the Netherlands), 1 µg/ml Hydrocortisone (H0135, Sigma-Aldrich), 1% Penicillin-Streptomycin (15140122, Gibco), and 10 ng/ml Epidermal Growth Factor (EGF) (E9644, Sigma-Aldrich). After thawing, HMEC-1 cells were cultured in T75 flasks for 3 days in a humidified incubator at 37 °C and 5% CO_2_.

### Microvessel-on-a-chip

The microvessel-on-a-chip model was created in the OrganoPlate^®^ 3-lane 40 (4003-400B, Mimetas B.V., Leiden, the Netherlands), a microfluidic titerplate with a 384-wells microtiter plate footprint and a bottom consisting of 40 microfluidic chips.^[62]^ As illustrated in Figure 1A and B, each glass bottomed chip consists out of three parallel microfluidic channels, separated by PhaseGuides^®^.^[63]^ The wells of the titerplate consist of standard virgin polystyrene and the chip bottom has a standard 1H coverslip thickness (150 µm). The top and bottom channels have a height of 220 µm and a width of 300 µm. The middle channel has a height of 220 µm and a width of 350 µm. Each channel has its own in- and outlet in separate wells which function as medium reservoirs and allow access to both sides of the channels.

To form the hollow tubular-shaped matrix in which the microvessel-on-a-chip was grown, a collagen I extracellular matrix gel was loaded in the middle inlet channel of each chip 1 day before cell seeding. The gel was prepared by mixing 5 mg/mL Cultrex Rat Collagen I matrix gel (3447-020-01, R&D systems, Bio-Techne, Minneapolis, MN, USA), 1 M HEPES (15630-056, Gibco), and 37 g/L HaHCO_3_ pH 9.5 (S5761, Sigma-Aldrich) in an 8:1:1 ratio to a final gel concentration of 4 mg/mL. The gel mixture was kept cold in an NEB cooler (T0225S, New Engeland BioLabs, Ipswich, MA, USA) during mixing and pipetting. Within 15 min after mixing, 2 µL of the gel was loaded into the OrganoPlate^®^. After loading, the gel was polymerized by incubating the OrganoPlate^®^ for 15 min in a humidified incubator at 37 °C and 5% CO_2_. After incubation, 20 µ L Dulbecco’s Phosphate-Buffered Saline (D-PBS) (14190094, Gibco) was added to all the middle channel inlets and the OrganoPlate^®^ was placed in a humidified incubator at 37 °C and 5% CO_2_ overnight. Due to the PhaseGuides^®^, the gel stayed contained into the middle channel and formed a meniscus towards the top and bottom channel.^[63]^

One day after gel loading, the D-PBS was removed from all the middle channel inlets and 50 µL of 28.5 µg/ml fibronectin solution (10 838 039 001; Roche Diagnostics Corp., Indianapolis, IN) in non-supplemented MCDB131 medium was added to all the top channel outlets. Next, the OrganoPlate^®^ was incubated for 30 min in a humidified incubator at 37 °C and 5% CO_2_. The HMEC-1 cells (passage number 24 or 26) were detached with trypsin/EDTA (CC-5012, Lonza), centrifuged at 220 *g*, 5 min, break 2 (Heraeus Biofuge, Thermo Scientific, Etten Leur, the Netherlands), and the cell pellet was resuspended into supplemented MCDB131 medium to reach a concentration of 1.0·10^7^ cells/mL. Subsequently, cell suspension was seeded into the channel by passive pumping, i.e. adding a 2µL droplet to the top channel inlet which will flow into the channel until surface tension between in and outlet is equilibrated ^[37]^. For the cell-free controls, 2 µ L of supplemented MCDB131 medium was added to the top channel outlet instead. After cell seeding, fibronectin solution was aspirated from the top channel outlet and 50 µ L of supplemented MCDB131 medium was added. To allow the cells to attach to the gel meniscus, the OrganoPlate^®^ was incubated under an angle of 75° on the MIMETAS plate stand in a humidified incubator at 37 °C and 5% CO_2_ for 2 h. After incubation, the plate was removed from the stand and 50 µL of supplemented MCDB131 medium was added to the top channel outlet. Next, the OrganoPlate^®^ was cultured under flow for 4 days by incubating the plate on the Mimetas OrganoFlow^®^ rocker (8 min intervals at 7 ° inclination angle) in a humidified incubator at 37 °C and 5% CO_2_. Medium from the top channel in- and outlets was refreshed 1 day after cell seeding. Experiments were performed 4 days after cell seeding. Before the experiment started, all middle and bottom channel in- and outlets of each microvessel were wetted by adding 50 µ L of supplemented MCDB131 medium to the, after which the plate was incubated for 5 min under a 7° inclination angle in a humidified incubator at 37 °C and 5% CO_2_.

### Gel Stiffness Measurements

The extracellular collagen I matrix gel was prepared as described above for three different gel Lot. numbers. Five gel drops of 10 µL were seeded in a 35 mm petridish. D-PBS was added in the same gel:D-PBS ratio of 1:10 as used in the OrganoPlate^®^ and the gel was incubated overnight in a humidified incubator at 37 °C and 5% CO_2_. Before measurements, the samples were fully submerged in D-PBS and the Young’s modulus of the gel was measured in nine different locations spaced 10 µm apart in a 3×3 grid using a Chiaro Nanoindenter (Optics11life, Amsterdam, the Netherlands) with a 3 µm tip radius probe (Optics11life) as previously reported.^[64]^

### Immunohistochemistry

Four days after cell seeding, immunohistochemistry was performed to visualize the α_v_β_3_ integrin expression on the HMEC-1 cells. All incubation steps were performed at room temperature and 100 µL was added to the top channel inlet and 50 µL was added to all the other channel in- and outlets, unless mentioned otherwise. First, the OrganoPlate^®^ was incubated with 4% paraformaldehyde (158127, Sigma-Aldrich) in D-PBS fixation solution under a 7° angle for 15 min. Next, the fixation solution was aspirated and chips were washed thrice by adding D-PBS and incubated under a 7° angle for 5 min. After washing, all chips were blocked by adding 5% goat serum (G6767, Sigma-Aldrich) in D-PBS blocking solution and incubating the OrganoPlate^®^ under a 7° angle for 30 min. Next, the blocking solution was aspirated and chips were incubated flat overnight at 4° C with 25 µ L biotinylated α_v_β_3_ antibody (1:50 dilution, 304412, BioLegend, San Diego, CA, USA) in the top channel in- and outlets and 15 µ L in the bottom channel in- and outlets. A mouse biotinylated IgG1 isotype control antibody (1:50 dilution, 2600520, Sony Biotechnology, San Jose, CA, USA) was used to assess the specificity of the α_v_β_3_ antibody. The next day, the antibody solution was aspirated and chips were washed thrice with 0.5% Tween-20 (P5927, Sigma-Aldrich) in D-PBS. After washing, all chips were blocked again using 5% goat serum in D-PBS under a 7° angle for 30 min. Then, chips were incubated in the dark with 25 µ L of anti-mouse Alexa Fluor 488 antibody (1:100 dilution, A-11029, Invitrogen, Thermo Fisher Scientific) in the top channel in- and outlets and 15 µL in the bottom channel in- and outlets under a 7° angle at room temperature for 1 h. Next, chips were washed thrice with 0.5% Tween-20 in D-PBS and 20 µg/mL Hoechst 33342 (H3570, Thermo Fisher Scientific) in D-PBS was added to all chips. After a 10 min incubation at room temperature in the dark under a 7° angle, the Hoechst 33342 solution was aspirated. Finally, 8 µL Vectashield (H-1400, Vector Laboratories Inc., Newark, CA, USA) was added to all channel inlets and incubated in the dark under a 7° angle for 15 min at room temperature. Thereafter, the plate was placed upside down and 3D z-stack images of 635 µm × 635 µm (512 × 512 pixels) with a z-step size of 0.8 µm were made using an A1R+ confocal microscope (Nikon Instruments, Amsterdam, the Netherlands) with a 20× water immersion objective (Cfi Apo LWD; numerical aperture 0.95; Nikon Instruments). Two laser channels and filter cubes were used: 1) Hoechst 33342 excited at 405 nm and detected at 450/50 nm (center wavelength/bandwidth), 2) Alexa Fluor 488 excited at 488 nm and detected at 525/50 nm.

### Microbubble Preparation and Targeting

Lipid-coated MBs with a C_4_F_10_ gas (F2 Chemicals, Preston, UK) core were produced using an indirect method with probe sonication for 1 min as previously described.^[65,66]^ Briefly, the lipids were dissolved in an organic solvent (chloroform:methanol 9:1 volume ratio) and mixed to obtain a molar ratio of 84.8% DSPC (Lipoid, Ludwigshafen, Germany), 8.2% PEG40-stearate (Sigma-Aldrich), 5.9% DSPE-PEG2000, and 1.1% biotinylated DSPE-PEG2000 (both from Avanti Polar Lipids, Alabaster, AL, USA). Lipid films were created by drying the lipids using argon gas (Linde Gas Benelux, Schiedam, the Netherlands) for ∼15 min and freeze drying for 2 h using an Alpha 1-2 LD plus Freeze dryer (Martin Christ GmbH, Osterode am Harz, Germany). Lipid films were rehydrated in 5 mL C_4_F_10_ saturated PBS and DiD (1,1’-dioctadecyl 3,3,3’,3’-tetramethylindodicarbocyanine perchlorate; D307, Thermo Fisher Scientific) lipid dye was added when fluorescent MBs were desired. The MBs were targeted to the α_v_β_3_ integrin using biotin-streptavidin bridging as previously described.^[55,67]^ Briefly, MBs were washed three to four times by centrifugation (400 *g*, 1 min, brake 9) using C_4_F_10_-saturated PBS until the subnatant was clear after which MBs were counted using a Coulter Counter Multisizer 3 (50 µm aperture tube, Beckman Coulter, Mijdrecht, the Netherlands). Next, 6·10^8^ MBs were incubated on ice for 30 min with 60 µg streptavidin (S4762, Sigma-Aldrich). Subsequently, streptavidin-conjugated MBs were washed once by centrifugation and incubated with 6 µg biotin anti-human α_v_β_3_ antibody (304412, BioLegend) on ice for 30 min. The α_v_β_3_-tMBs were washed by centrifugation for one more time and the final α_v_β_3_-tMB concentration was determined by counting using the Coulter Counter Multisizer 3.

To add the α_v_β_3_-tMBs to the microvessels, they were diluted to a concentration of 1.5·10^7^ α_v_β_3_-tMBs/mL in 37 °C supplemented MCDB131 medium. Then, the plate was placed under an angle of 75° on the MIMETAS plate stand in such a way that the gel-cell interface layer was the highest point of the microvessel. While the plate was on the MIMETAS plate stand, 40 µ L of the diluted α_v_β_3_-tMBs solution was added to the top channel inlet and 30 µ L to the top channel outlet, followed by a 5 min incubation step to allow the α_v_β_3_-tMBs to bind to the cells in contact with the gel.

### Experimental Setup

A custom-built Nikon A1R+ confocal microscopic setup was used to image the OrganoPlate^®^ (**Figure 10**A). This upright microscope was equipped with a DS-Fi3 (Nikon Instruments) color camera for brightfield and widefield fluorescence imaging, brightfield light source (Nikon Instruments) and metal halide light source (Fiber Illuminator Intensilight; Nikon Instruments). A V-shaped 37 °C water bath was placed underneath the microscope to allow for simultaneous temperature control of the OrganoPlate^®^, microscopic imaging and ultrasound insonification. A single element focused transducer (2.25 MHz center frequency; 76.2 mm focal length; -6 dB beam width at 2 MHz of 3 mm; V305; Panametrics-NDT, Olympus, Waltham, MA, USA) was positioned in the water bath under an incidence angle of 45° in relation to the OrganoPlate^®^. The pressure output from the transducer was previously calibrated using a 1 mm needle hydrophone (Precision Acoustic, Dorchester, UK). The ultrasound focus was aligned to correspond with the optical focus of the microscope as previously described.^[68]^ The 2 MHz frequency ultrasound waves were generated using an arbitrary waveform generator (33220A 20MHz function, Agilent, Palo Alto, CA, USA) and amplified 50 dB (ENI 2100L, Electronics & Innovation, Rochester, NY, USA) to generate 100-850 kPa PNP in water and 90-750 kPa PNP *in situ* in the chips in the OrganoPlate^®^.^[30]^ Ultrasound was applied in 10 separate bursts of 10 or 1000 cycles per burst, which were applied every 3 s for a total of 30 s.

**Figure 10.**
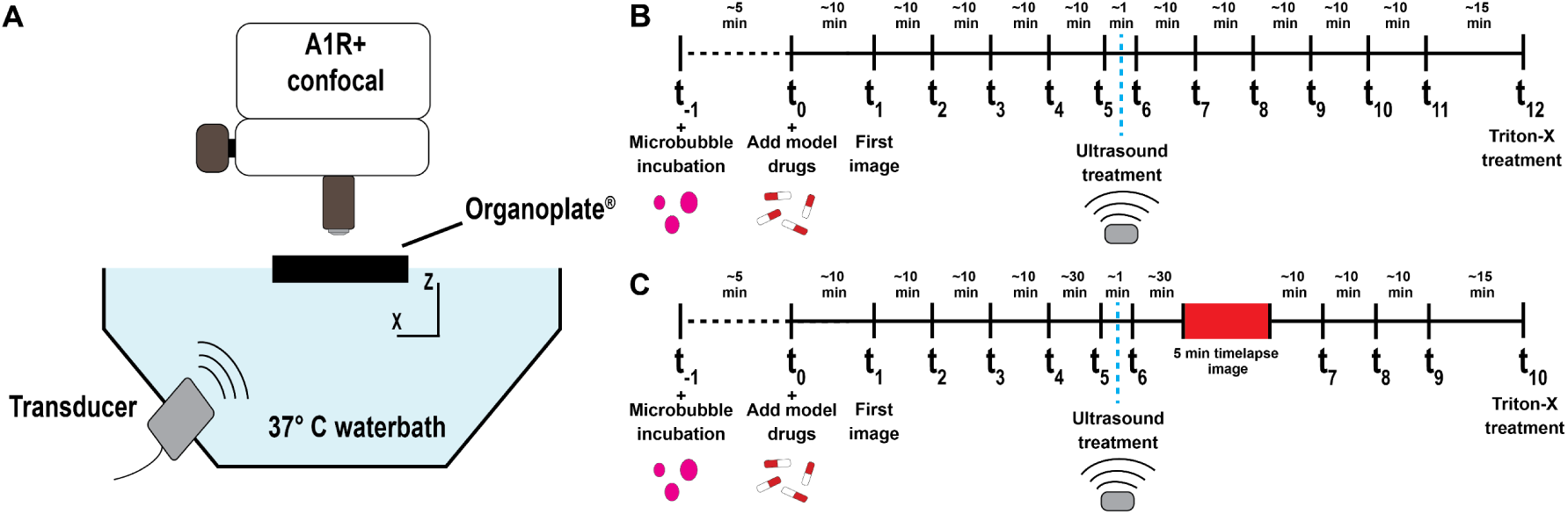
Imaging setup and experimental timeline for the barrier integrity (BI) assay. A) The optical imaging system allows for brightfield and epi-fluorescent microscopy imaging with a DS-Fi3 color camera (Nikon Instruments) and confocal microscopy imaging with the A1R+ scan head (Nikon Instruments). Ultrasound was applied under an angle of 45° in a 37 °C water bath underneath the OrganoPlate^®^. B) Experimental timeline of the BI assay for all treatment conditions. C) Experimental timeline of BI assay including 5-min timelapse imaging directly following treatment on a subset of treatment conditions. For both timelines, the α_V_β_3_-targeted microbubbles were added at t_-1_ and the assays were started by adding the BI solution containing the model drugs at t_0_.

### Barrier Integrity Assay

To assess the permeability of a microvessel-on-a-chip before and after different treatments, a barrier integrity (BI) assay was performed. The timeline of the experiment is shown in Figure 10B. Before the BI assay, the plate was divided into two sections and both sections were prepared and imaged separately to be able to achieve the desired imaging speed. Following the 5 min α_v_β_3_-tMB incubation (t_-1_; see *Microbubble Preparation and Targeting*) in the vessel channel in the first section of the plate, all channels were aspirated and the middle and bottom channels were wetted by adding 20 µL supplemented MCDB131 medium to the channel in- and outlet. A BI solution containing model drugs was prepared consisting of 0.5 mg/mL 150 kDa FITC dextran (46946, Sigma-Aldrich) and 25 µg/mL propidium iodide (PI) (P4864, Sigma-Aldrich) in 37 °C supplemented MCDB131 medium. The BI assay was started by adding 40 µ L BI solution to the top channel inlet and 30 µ L to the top channel outlet (t_0_) and placing the Organoplate^®^ in the 37 °C water bath (Figure 10A) with the inlet openings to the top. Using a 4× air objective (Cfi Plan Fluor, 17.2 mm working distance, Nikon Instruments), sequences of one brightfield and two fluorescent widefield images of 2880 µm × 2048 µm (2880 × 2048 pixels) were recorded ∼10 min apart with the DS-Fi3 color camera. The fluorescent BI solution was detected using the following excitation (Ex), dichroic mirror (DM), and emission (Em) filters: for FITC-dextran Ex469/35, DM497 and Em525/39; for PI Ex542/20, DM570, and Em620/52 (center wavelength/bandwidth in nanometers; Semrock Inc., Rochester, NY, USA). After adding the BI solution, the initial state of the microvessels was imaged for at least 30 min using five images (t_1_-t_5_), including an image just before treatment (t_5_), to assess the permeability and quality of the microvessels. After the initial state, different treatments, depending on the conditions, were randomly applied for 30 s during which a brightfield video was made (∼2.5 frames per sec (fps)). In total 14 treatment conditions were investigated: ten different ultrasound plus α_v_β_3_-tMB treatments and four controls (sham; tMBs only; 750 kPa 10×10 cycles ultrasound only; 750 kPa 10×1000 cycles ultrasound only). In addition, cell free chips were included which contained no cells and only ECM gel. Directly after each treatment, another sequence of images was made (t_6_). The sequence of imaging was continued for five more time points ∼10 min apart (t_7_-t_11_) for ∼45 min. Next, the barrier was disrupted by killing the cells to ensure that the observed permeability was a result of the cellular layer and not an artefact induced by for example liquid flow or the gel. To kill cells, Triton-X (T8787, Sigma-Aldrich) at a final concentration of 0.1% was added to all top channel inlets followed by one final sequence of images (t_12_). The above procedures were repeated for the second section of the plate starting at t_-1_. In addition, a BI assay with a 5-min timelapse in brightfield and fluorescence directly following treatment was recorded (0.2 fps) for a subset of treatment conditions, see timeline in Figure 10C. In the initial state, i.e. up to t_5_, and barrier disrupting step by killing cells (t_10_), all procedures were identical as described above. However, here t_6_ was the sequence of images directly before treatment. Directly after the 5-min timelapse, the sequence of imaging was continued for three more time points ∼10 min apart (t_7_-t_9_) for ∼45 min.

### Live Cell 3D Imaging

Four days after cell seeding, live cell imaging was performed to quantify the number of bound α_v_β_3_-tMBs in the microvessel and visualize sonoporation. The Organoplate^®^ was wetted and α_v_β_3_-tMBs were incubated as described above (see *Microbubble Preparation and Targeting*). Staining solution with a final concentration of 20 µg/mL Hoechst 33342 (H3570, Thermo Fisher Scientific), 12 µg/mL CellMask^TM^ Green Plasma Membrane Stain (C37608, Thermo Fisher Scientific) and 25 µg/mL Propidium Iodide (PI) was prepared in 37 °C supplemented MCDB131 medium. Next, staining solution was added as for the BI solution described above in the BI assay section. Following incubation, the plate was imaged upside down such that the microvessels were within the working distance of the Cfi Apo LWD 20× water immersion objective (Nikon Instruments). Sixteen 3D z-stack images of 635 µm × 635 µm (512 × 512 pixels) with a z-step size of 0.8 µm were made in eight different microvessels using an A1R+ confocal microscope (Nikon Instruments). Four laser channels and filter cubes were used: 1) Hoechst 44332 excited at 405 nm and detected at 450/50 nm (center wavelength/bandwidth), 2) CellMask^TM^ Green Plasma Membrane Stain excited at 488 nm and detected at 525/50 nm, 3) PI excited at 561 nm and detected at 595/50 nm, 4) DiD excited at 640 nm and detected at 700/75 nm. Channel 1 and 4 were excited and detected simultaneously because there is no spectral overlap between Hoechst 44332 and DiD. Next, the Organoplate® was placed back into the original position with the bottom of the plate inside the water bath and ultrasound was applied as described above. After treatment, the Organoplate^®^ was positioned upside down again and the microvessels were imaged once more which was 14-70 min after treatment.

### Cell Viability Protocol

To investigate whether the treatment conditions affected cell viability and eliminate that PI uptake was due to cell death, the viability of the cells within the entire microvessel was assessed using a WST-8 chromatography assay. The BI assay up to timepoint t_6_ was performed as described above and illustrated in Figure 10B. At timepoint t_6_ of the BI assay, the Organoplate^®^ was taken out of the imaging setup and 0.1% Triton-X (final concentration; T8787, Sigma-Aldrich) was added to the desired chips and incubated for 10 min at 37° C. Next, all chips were aspirated and 25 µ L of WST-8 working solution was added to the top and bottom channel in- and outlets. The WST-8 working solution was made by diluting the WST-8 stock solution (96992, Sigma-Aldrich) 1:11 in supplemented MCDB131 medium. Next, the Organoplate^®^ was incubated for 30 min on the OrganoFlow^®^ rocker as described above. This was followed by placing the Organoplate^®^ flat for 5 min to stop the flow. Subsequently, the absorbance at a wavelength of 450 nm was measured in the in- and outlets of the top channel using a spectrophotometer (Spectramax iD3, Molecular Devices, LLC, San Jose, CA, USA). The average time elapsed between the last treatment and the absorbance reading was 78 min (range 73-83 min). The negative controls were: MCDB131 medium without WST-8 reagent and three cell-free conditions with WST-8 reagent of which one was treated with α_v_β_3_-tMBs only and one with ultrasound (750 kPa 10×1000 cycles) plus α_v_β_3_-tMBs. The positive control was the microvessels incubated with 0.1% Triton-X as described above, which kills all the cells.

### Bound α_v_β_3_-tMB Quantification Analysi

Live cell 3D stack images were divided into three segments: 1) the bottom of the microvessel, i.e., the 2D cell layer at the bottom of the Organoplate_®_ up to the top of the phaseguide; 2) the gel-cell interface; and 3) the top of the microvessel, i.e., the 2D cell layer on the top of the microvessel. Particles were counted using the ‘Small particles geometry’ analysis in the ‘Object Analyzer’ tool, combined with a watershed segmentation of 10%, in the Huygens Professional image analysis software (Scientific Volume Imaging, Hilversum, the Netherlands). Using the Coulter Counter Multisizer 3 measurements, particles were classified as debris when the particle volume was below the *d*10%, (i.e., the α_v_β_3_-tMB diameter below which 10% of the cumulative amount of α_v_β_3_-tMBs was found), classified as one α_v_β_3_-tMB when the particle volume was between the *d*10% and *d*90% (i.e., the tMB diameter below which 90% of the cumulative amount of α_v_β_3_-tMBs was found) or classified as multiple α_v_β_3_-tMBs when the particle volume was above the *d*90%. When classified as multiple α_v_β_3_-tMBs, the particle volume was divided by the mean voxel volume (i.e. 18 voxels) of one α_v_β_3_-tMB (i.e., mean α_v_β_3_-tMB diameter) and all decimal values were rounded up to calculate the number of single α_v_β_3_-tMBs in the multiple particle.

### Data Analysis BI Assay

All data analysis was performed using MATLAB R2021b (The MathWorks Inc., Natick, MA, USA). Leakage was quantified by drawing a region of interest (ROI) in the top microvessel and middle gel channels and measuring the average pixel intensity of the green FITC-dextran signal per ROI. The leakage percentage over time was calculated by dividing the fluorescent intensity of the middle gel channel ROI (*I_gel_*) by the intensity of the top microvessel channel ROI (*I_ves_*):

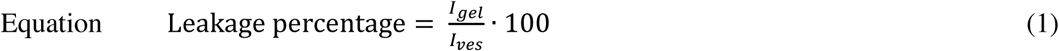

If the leakage value exceeded 50% within the first 30 min of the BI assay, the microvessels were deemed unsuitable for the assay and excluded. The Apparent Permeability (P_app_) of the vessels was calculated in the initial state (i.e., before treatment) and after treatment using the Excel solver tool provided by Mimetas.^[69]^ For the BI assay with the 5-min timelapse (as illustrated in Figure 10C), the P_app_ was also calculated for the 5-min period directly following the ultrasound treatment.

### Data analysis sonoporation

All data analysis was performed using MATLAB R2021b (The MathWorks Inc.). Sonoporation was quantified by drawing a ROI on the gel-cell interface part of the top microvessel channel and measuring the average PI signal intensity per ROI. Percentage increase was calculated using the difference in signal in the last image before (t_5_) and the first image after (t_6_) treatment. Image t_6_ was always taken within 45 s after the last ultrasound pulse. N.B. These time points relate to the timeline illustrated in Figure 10B.

### Data Analysis

*α_v_β_3_-tMB Displacement, α_v_β_3_-tMB Clustering and Relation with Vascular Permeability/PI Uptake in vessel segments:* α_v_β_3_-tMB displacement and clustering were extracted from the brightfield videos recorded during ultrasound treatment and analyzed with MATLAB R2019a (The MathWorks Inc.). First, the videos were denoised using a Gaussian filter, and cropped so only the top microvessel channel was imaged. Second, the difference in pixel intensity before and after each ultrasound pulse was analyzed. Since α_v_β_3_-tMB scatter light and scattering effectively decreases the light that reaches the sensor, α_v_β_3_-tMB displacement will lead to a pixel intensity increase at the position where the α_v_β_3_-tMB displaced from. Therefore, α_v_β_3_-tMB displacement was quantified as the cumulative pixel intensity increase between frames. A α_v_β_3_-tMB cluster was defined as a group of at least 40 pixels (40 µm^2^, i.e., 3.3 times the mean α_v_β_3_-tMB area) with an intensity decrease, discarding those intersecting the boundaries of the cropped vessel. To reject noise in both the pixel intensity increase and decrease, only intensity changes above a heuristic threshold determined from the control recordings were considered. The distance between α_v_β_3_-tMB clusters was obtained using the MATLAB built-in *pdist* function.

The relation between α_v_β_3_-MB clusters and α_v_β_3_-MB displacement, vascular permeability increase, and PI increase was locally assessed. For this, the recording of the fluorescent channels before and after ultrasound treatment were loaded into MATLAB R2021b (The MathWorks Inc.) and filtered with a 2D wiener filter. The vascular permeability increase was measured on the middle gel channel, whereas PI increase was measured on the top microvessel channel. In order to obtain spatial information, 200 µm wide segments were defined along the x-direction of the ROI (see red-dotted rectangles in Figure 6A), measuring the number of α_v_β_3_-tMB clusters, cumulative α_v_β_3_-tMB displacement, vascular permeability increase, and PI increase in each segment. Increases in vascular permeability and PI were only considered relevant if they exceeded a threshold, i.e., 0.01 A.U. for local leakage and 0.005 A.U. for local PI increase, which was defined as the quantile 96 value from the segments belonging to the control-treated microvessels (**Figure S8**).

### Statistics

All statistical analysis was performed in SPSS (IBM, Armonk, NY, USA). The Shapiro-Wilk test of normality was used to assess whether data were parametric or non-parametric. For non-parametric data, the Mann-Whitney U test was used while parametric data was tested for equality of variances using the Levene’s test after which a student’s t-test was used for data with equal variances or the Welch’s t-test for data with unequal variances. Reported p-values are two-tailed and p < 0.05 is indicated with *, p < 0.01 is indicated with **, and p < 0.001 is indicated with ***. In all Figures, except for Figure 9, the indicated significance represents changes compared to the four control conditions (i.e., sham, α_v_β_3_-tMBs only, and ultrasound only (750 kPa 10×10 cycles and 750 kPa and 10×1000 cycles)) and the differences within these significant groups. In Figure 9, the indicated significance represents the differences compared to the one positive and four negative control conditions. Pearsons’s correlation was performed on all data points above the control threshold in Figure 8 using Graphpad Prism 9 (Dotmatics, Boston, MA, USA).

## Supporting Information

Supporting Information is available from the Wiley Online Library or from the author.

## Acknowledgements

This work was funded in majority by the Applied and Engineering Sciences (TTW) (Vidi-project 17543), part of the Dutch Research Council NWO, awarded to K.K. J.C. and G.H.K. gratefully acknowledge financial support from the Dutch Research Council NWO (project number VI.C.182.004 of the NWO Talent Programme).

The authors would like to thank Robert Beurskens, Phoei Ying Tang and Cheryl Mok from the Biomedical Engineering of the Department of Cardiology, Michiel Manten from the Department of Experimental Medical Instrumentation and Marcel Sluijter from the Laboratory of Paediatrics, all from Erasmus MC, the Netherlands, for technical assistance.

## Conflict of Interest

Kristina Bishard and Sebastiaan J. Trietsch are employees of MIMETAS BV, which is marketing the OrganoPlate and Sebastiaan J. Trietsch is a shareholder of MIMETAS BV. OrganoPlate is a registered trademark of MIMETAS BV. The other authors declare no conflict of interest.

## Data Availability Statement

The data that support the findings of this study are available from the corresponding author upon reasonable request.

## Table of contents

To gain mechanistic insight into vascular permeability increases by ultrasound-activated microbubbles, accurate models are needed. Here, microvessels-on-chip are presented with a membrane-free extravascular space as original method. The onset rate and amount of vascular permeability increase shows distinct and significant differences between 10 and 1000 cycle-treatments of ultrasound (2 MHz, 750 kPa) with α_v_β_3_-targeted microbubbles, thereby further unraveling the mechanism.

Bram Meijlink, Gonzalo Collado Lara, Kristina Bishard, James P. Conboy, Simone A.G. Langeveld, Gijsje H. Koenderink, Antonius F.W. van der Steen, Nico de Jong, Inés Beekers, Sebastiaan J. Trietsch, Klazina Kooiman

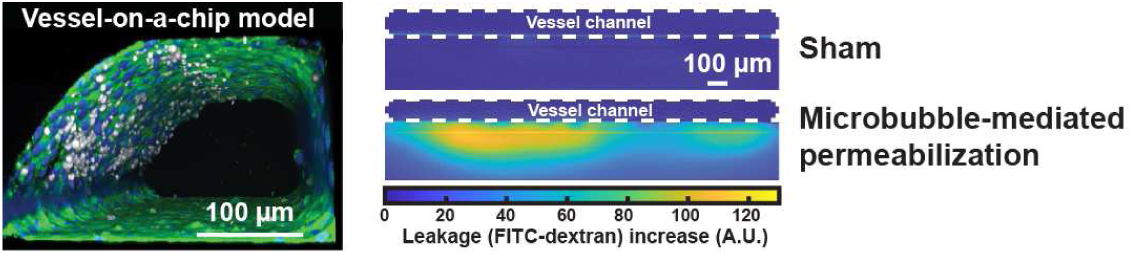
Characterizing microbubble-mediated permeabilization in a vessel-on-a-chip model.

## References

[1] A. A. Van der Veldt, M. Lubberink, I. Bahce, M. Walraven, M. P. de Boer, H. N. Greuter, N. H. Hendrikse, J. Eriksson, A. D. Windhorst, P. E. Postmus, H. M. Verheul, E. H. Serne, A. A. Lammertsma, E. F. Smit, Cancer Cell 2012, 21, 82.

[2] S. Heskamp, O. C. Boerman, J. D. Molkenboer-Kuenen, W. J. Oyen, W. T. van der Graaf, H. W. van Laarhoven, Int J Cancer 2013, 133, 307.

[3] M. Arjaans, T. H. Oude Munnink, S. F. Oosting, A. G. Terwisscha van Scheltinga, J. A. Gietema, E. T. Garbacik, H. Timmer-Bosscha, M. N. Lub-de Hooge, C. P. Schroder, E. G. de Vries, Cancer Res 2013, 73, 3347.

[4] K. Kooiman, S. Roovers, S. A. G. Langeveld, R. T. Kleven, H. Dewitte, M. A. O’Reilly, J. M. Escoffre, A. Bouakaz, M. D. Verweij, K. Hynynen, I. Lentacker, E. Stride, C. K. Holland, Ultrasound Med Biol 2020, 46, 1296.

[5] J. Deprez, G. Lajoinie, Y. Engelen, S. C. De Smedt, I. Lentacker, Adv Drug Deliv Rev 2021, 172, 9.

[6] S. R. Wilson, P. N. Burns, Y. Kono, Ultrasound Med Biol 2020, 46, 1059.

[7] T. R. Porter, S. B. Feinstein, F. J. Ten Cate, A. E. van den Bosch, Ultrasound Med Biol 2020, 46, 1071.

[8] B. Helfield, X. Chen, S. C. Watkins, F. S. Villanueva, Proc Natl Acad Sci U S A 2016, 113, 9983.

[9] I. Beekers, M. Vegter, K. R. Lattwein, F. Mastik, R. Beurskens, A. F. W. van der Steen, N. de Jong, M. D. Verweij, K. Kooiman, Journal of Controlled Release 2020, 322, 426.

[10] B. D. Meijering, L. J. Juffermans, A. van Wamel, R. H. Henning, I. S. Zuhorn, M. Emmer, A. M. Versteilen, W. J. Paulus, W. H. van Gilst, K. Kooiman, N. de Jong, R. J. Musters, L. E. Deelman, O. Kamp, Circ Res 2009, 104, 679.

[11] K. Kooiman, H. J. Vos, M. Versluis, N. de Jong, Adv. Drug Deliver. Rev. 2014, 72C, 28.

[12] J. Park, Z. Fan, C. X. Deng, J Biomech 2011, 44, 164.

[13] R. E. Kumon, M. Aehle, D. Sabens, P. Parikh, Y. W. Han, D. Kourennyi, C. X. Deng, Ultrasound Med Biol 2009, 35, 494.

[14] Z. Fan, H. Liu, M. Mayer, C. X. Deng, Proc Natl Acad Sci U S A 2012, 109, 16486.

[15] I. Beekers, F. Mastik, R. Beurskens, P. Y. Tang, M. Vegter, A. F. W. van der Steen, N. de Jong, M. D. Verweij, K. Kooiman, Ultrasound in Medicine & Biology 2020, 46, 2017.

[16] I. Beekers, S. A. G. Langeveld, B. Meijlink, A. F. W. van der Steen, N. de Jong, M. D. Verweij, K. Kooiman, J Control Release 2022, 347, 460.

[17] B. Helfield, X. Chen, S. C. Watkins, F. S. Villanueva, Ultrasound Med Biol 2020, 46, 1686.

[18] S. M. Stieger, C. F. Caskey, R. H. Adamson, S. Qin, F. R. Curry, E. R. Wisner, K. W. Ferrara, Radiology 2007, 243, 112.

[19] F. Vlachos, Y. S. Tung, E. Konofagou, Magn Reson Med 2011, 66, 821.

[20] P. T. Yemane, A. K. O. Aslund, S. Snipstad, A. Bjorkoy, K. Grendstad, S. Berg, Y. Morch, S. H. Torp, R. Hansen, C. L. Davies, Ultrasound Med Biol 2019, 45, 3028.

[21] G. Dimcevski, S. Kotopoulis, T. Bjanes, D. Hoem, J. Schjott, B. T. Gjertsen, M. Biermann, A. Molven, H. Sorbye, E. McCormack, M. Postema, O. H. Gilja, J Control Release 2016, 243, 172.

[22] S. Roovers, T. Segers, G. Lajoinie, J. Deprez, M. Versluis, S. C. De Smedt, I. Lentacker, Langmuir 2019.

[23] G. Lajoinie, I. De Cock, C. C. Coussios, I. Lentacker, S. Le Gac, E. Stride, M. Versluis, Biomicrofluidics 2016, 10, 011501.

[24] E. Memari, F. Hui, H. Yusefi, B. Helfield, J Control Release 2023, 358, 333.

[25] A. Pollet, J. M. J. den Toonder, Bioengineering (Basel) 2020, 7.

[26] E. K. Juang, I. De Cock, C. Keravnou, M. K. Gallagher, S. B. Keller, Y. Zheng, M. Averkiou, Langmuir 2019, 35, 10128.

[27] G. Silvani, C. Scognamiglio, D. Caprini, L. Marino, M. Chinappi, G. Sinibaldi, G. Peruzzi, M. F. Kiani, C. M. Casciola, Small 2019, 15, e1905375.

[28] S. J. Trietsch, E. Naumovska, D. Kurek, M. C. Setyawati, M. K. Vormann, K. J. Wilschut, H. L. Lanz, A. Nicolas, C. P. Ng, J. Joore, S. Kustermann, A. Roth, T. Hankemeier, A. Moisan, P. Vulto, Nat Commun 2017, 8, 262.

[29] L. de Haan, J. Suijker, R. van Roey, N. Berges, E. Petrova, K. Queiroz, W. Strijker, T. Olivier, O. Poeschke, S. Garg, L. J. van den Broek, Int J Mol Sci 2021, 22, 8234.

[30] I. Beekers, T. van Rooij, M. D. Verweij, M. Versluis, N. de Jong, S. J. Trietsch, K. Kooiman, IEEE Trans Ultrason Ferroelectr Freq Control 2018, 65, 570.

[31] T. van Rooij, I. Skachkov, I. Beekers, K. R. Lattwein, J. D. Voorneveld, T. J. A. Kokhuis, D. Bera, Y. Luan, A. F. W. van der Steen, N. de Jong, K. Kooiman, J Control Release 2016, 238, 197.

[32] K. Kooiman, M. Emmer, M. Foppen-Harteveld, A. van Wamel, N. de Jong, IEEE Trans Biomed Eng 2010, 57, 29.

[33] D. Zeng, T. Juzkiw, A. T. Read, D. W. Chan, M. R. Glucksberg, C. R. Ethier, M. Johnson, Biomech Model Mechanobiol 2010, 9, 19.

[34] N. Fekete, A. V. Beland, K. Campbell, S. L. Clark, C. A. Hoesli, Transfusion 2018, 58, 1800.

[35] J. Liu, H. Zheng, P. S. Poh, H. G. Machens, A. F. Schilling, Int J Mol Sci 2015, 16, 15997.

[36] K. E. Sung, G. Su, C. Pehlke, S. M. Trier, K. W. Eliceiri, P. J. Keely, A. Friedl, D. J. Beebe, Biomaterials 2009, 30, 4833.

[37] V. van Duinen, W. Stam, V. Borgdorff, A. Reijerkerk, V. Orlova, P. Vulto, T. Hankemeier, A. J. van Zonneveld, J Vis Exp 2019.

[38] A. G. Monteduro, S. Rizzato, G. Caragnano, A. Trapani, G. Giannelli, G. Maruccio, Biosens Bioelectron 2023, 231, 115271.

[39] V. van Duinen, A. van den Heuvel, S. J. Trietsch, H. L. Lanz, J. M. van Gils, A. J. van Zonneveld, P. Vulto, T. Hankemeier, Sci Rep 2017, 7, 18071.

[40] A. Nicolas, F. Schavemaker, K. Kosim, D. Kurek, M. Haarmans, M. Bulst, K. Lee, S. Wegner, T. Hankemeier, J. Joore, K. Domansky, H. L. Lanz, P. Vulto, S. J. Trietsch, Lab Chip 2021, 21, 1676.

[41] X. Zhao, C. Pellow, D. E. Goertz, Theranostics 2023, 13, 250.

[42] T. Nhan, A. Burgess, E. E. Cho, B. Stefanovic, L. Lilge, K. Hynynen, Journal of Controlled Release 2013, 172, 274.

[43] D. H. Thomas, V. Sboros, M. Emmer, H. Vos, N. de Jong, IEEE Trans Ultrason Ferroelectr Freq Control 2013, 60, 105.

[44] C. F. Caskey, S. M. Stieger, S. Qin, P. A. Dayton, K. W. Ferrara, J Acoust Soc Am 2007, 122, 1191.

[45] T. Frey, Cytometry 1995, 21, 265.

[46] W. D. Wilson, C. R. Krishnamoorthy, Y. H. Wang, J. C. Smith, Biopolymers 1985, 24, 1941.

[47] Y. Hu, J. M. Wan, A. C. Yu, Ultrasound Med Biol 2013, 39, 2393.

[48] I. De Cock, E. Zagato, K. Braeckmans, Y. Luan, N. de Jong, S. C. De Smedt, I. Lentacker, J Control Release 2015, 197, 20.

[49] T. J. Kokhuis, V. Garbin, K. Kooiman, B. A. Naaijkens, L. J. Juffermans, O. Kamp, A. F. van der Steen, M. Versluis, N. de Jong, Ultrasound Med Biol 2013, 39, 490.

[50] V. Garbin, M. Overvelde, B. Dollet, N. de Jong, D. Lohse, M. Versluis, Phys Med Biol 2011, 56, 6161.

[51] X. Chen, J. Wang, J. J. Pacella, F. S. Villanueva, Ultrasound Med Biol 2016, 42, 528.

[52] A. A. Doinikov, Research Signpos 2005.

[53] S. Kotopoulis, M. Postema, Ultrasonics 2010, 50, 260.

[54] C. Lazarus, A. N. Pouliopoulos, M. Tinguely, V. Garbin, J. J. Choi, J Acoust Soc Am 2017, 142, 3135.

[55] I. Skachkov, Y. Luan, A. F. van der Steen, N. de Jong, K. Kooiman, IEEE Trans Ultrason Ferroelectr Freq Control 2014, 61, 1661.

[56] T. J. A. Kokhuis, B. A. Naaijkens, L. J. M. Juffermans, O. Kamp, A. F. W. van der Steen, M. Versluis, N. de Jong, Applied Physics Letters 2017, 111.

[57] C. Pellow, M. A. O’Reilly, K. Hynynen, G. Zheng, D. E. Goertz, Nano Lett 2020, 20, 4512.

[58] D. Gonzalez-Rodriguez, C. Morel, E. Lemichez, Adv Exp Med Biol 2020, 1267, 101.

[59] L. Claesson-Welsh, E. Dejana, D. M. McDonald, Trends Mol Med 2021, 27, 314.

[60] J. T. Belcik, B. P. Davidson, A. Xie, M. D. Wu, M. Yadava, Y. Qi, S. Liang, C. R. Chon, A. Y. Ammi, J. Field, L. Harmann, W. M. Chilian, J. Linden, J. R. Lindner, Circulation 2017, 135, 1240.

[61] N. Rong, M. Zhang, Y. Wang, H. Wu, H. Qi, X. Fu, D. Li, C. Yang, Y. Wang, Z. Fan, Ultrason Sonochem 2020, 67, 105125.

[62] S. J. Trietsch, G. D. Israels, J. Joore, T. Hankemeier, P. Vulto, Lab Chip 2013, 13, 3548.

[63] P. Vulto, S. Podszun, P. Meyer, C. Hermann, A. Manz, G. A. Urban, Lab Chip 2011, 11, 1596.

[64] N. L. G. Mattei, E.J. Breel, Optics11 White Paper 2017.

[65] S. A. G. Langeveld, C. Schwieger, I. Beekers, J. Blaffert, T. van Rooij, A. Blume, K. Kooiman, Langmuir 2020, 36, 3221.

[66] S. A. G. Langeveld, B. Meijlink, I. Beekers, M. Olthof, A. F. W. van der Steen, N. de Jong, K. Kooiman, Pharmaceutics 2022, 14, 311.

[67] K. Kooiman, M. Foppen-Harteveld, A. F. van der Steen, N. de Jong, J Control Release 2011, 154, 35.

[68] I. Beekers, K. R. Lattwein, J. J. P. Kouijzer, S. A. G. Langeveld, M. Vegter, R. Beurskens, F. Mastik, R. Verduyn Lunel, E. Verver, A. F. W. van der Steen, N. de Jong, K. Kooiman, Ultrasound Med Biol 2019, 45, 2575.

[69] C. Soragni, T. Vergroesen, N. Hettema, G. Rabussier, H. L. Lanz, S. J. Trietsch, L. J. de Windt, C. P. Ng, STAR Protoc 2023, 4, 102051.

