## Supporting Information for "Characterizing microbubble-mediated permeabilization in a vessel-on-a-chip model"

**Video S1.** Three-dimensional confocal microscopy recording animation of the microvessel shown in Figure 1B. Staining was performed for cell nuclei (pseudo-colored in blue), cell membranes (pseudo-colored in green), sonoporated/dead cells (pseudo-colored in red) and $\alpha_v\beta_3$-targeted microbubbles (pseudo-colored in white).

**Video S2.** Spatial distribution of the leakage increase during the 5-min time-lapse imaging after treatment (corresponding to Figure 4D). Time in min:s; scalebar represents 500 µm.

**Video S3.** Representative example video illustrating the $\alpha_v\beta_3$-targeted microbubble displacement during the 750 kPa 10×10 cycles ultrasound treatment (corresponding to Figure 6A). Time in s; scalebar represents 500 µm.

**Video S4.** Representative example video illustrating the $\alpha_v\beta_3$-targeted microbubble displacement and clustering during the 750 kPa 10×1000 cycles ultrasound treatment (corresponding to Figure 6B). Time in s; scalebar represents 500 µm.

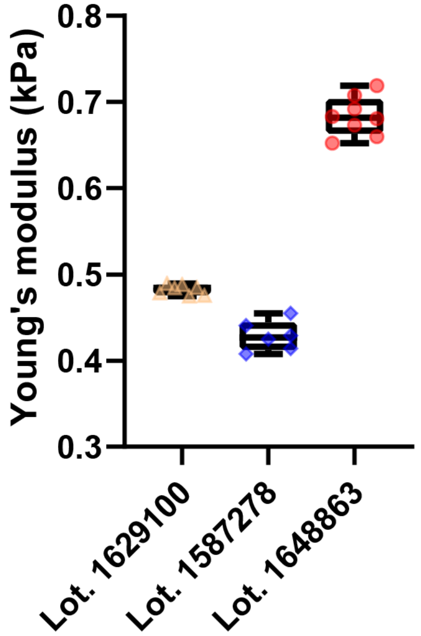


**Figure S1.** Gel stiffness measurements of three different Lot. Numbers of the collagen I extracellular matrix gel. Boxplot represent the median and the boxes indicate the 25$^{th}$ and 75$^{th}$ percentiles with whiskers ranging from the minimum to maximum value.

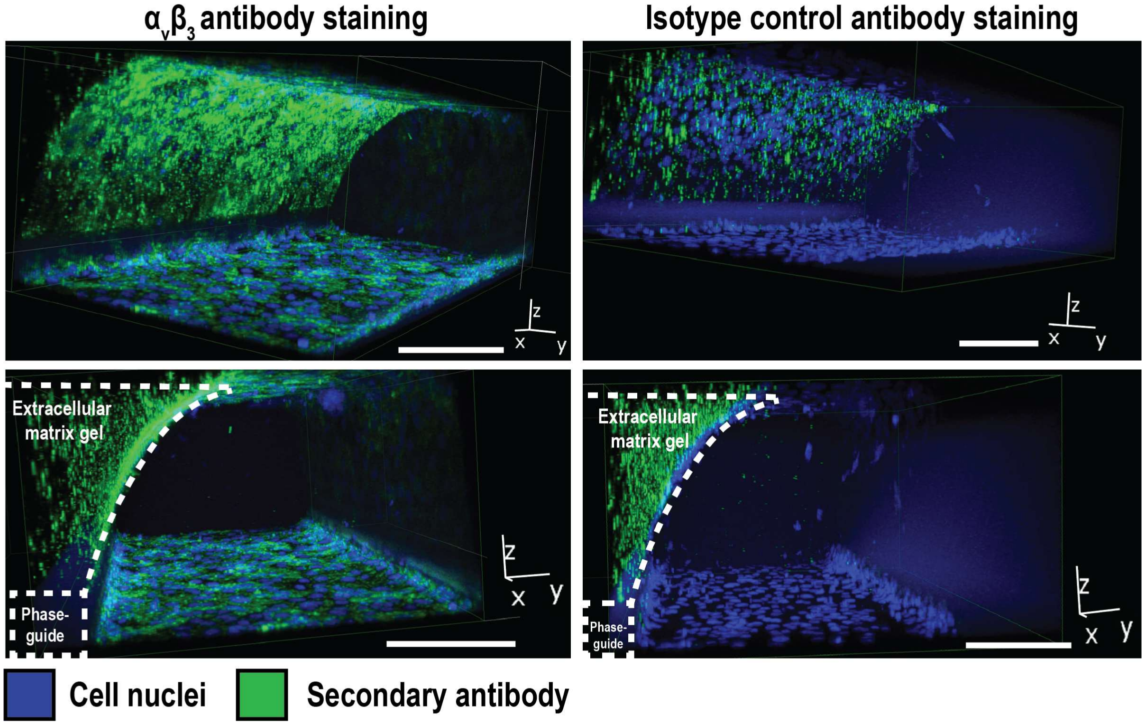


**Figure S2**. Three-dimensional immunofluorescence images of fixated microvessels-on-chip stained for $\alpha_v\beta_3$ integrin (left) and isotype control antibodies (right). The top and bottom images are two different views from the same part of the microvessel. The scalebar represents 100 µm for y-z direction; x-direction is 635 µm.

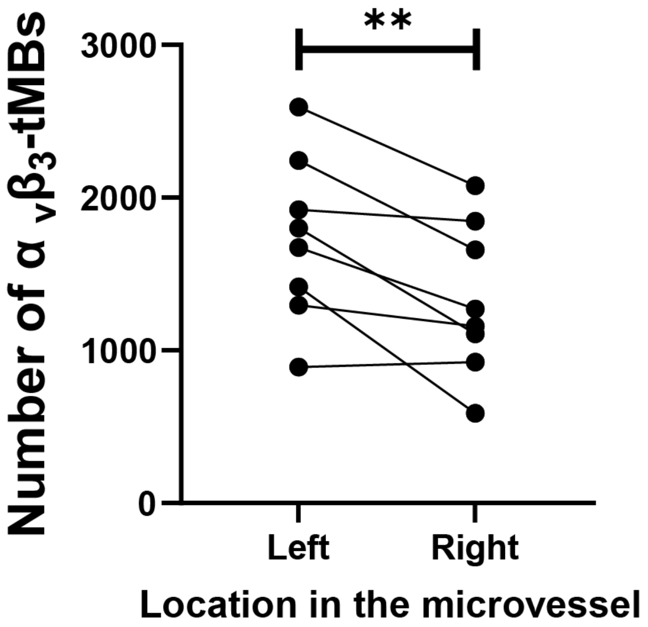


**Figure S3.** The number of bound $\alpha_v\beta_3$-tMBs on the gel-cell interface compared between the left (indicated with locations I in Figure 1A) and the right (indicated with locations II in Figure 1A) side of the same vessels. Statistical significance is indicated with **$p<0.01$ $\alpha_v\beta_3$-tMBs = $\alpha_v\beta_3$-targeted microbubbles.

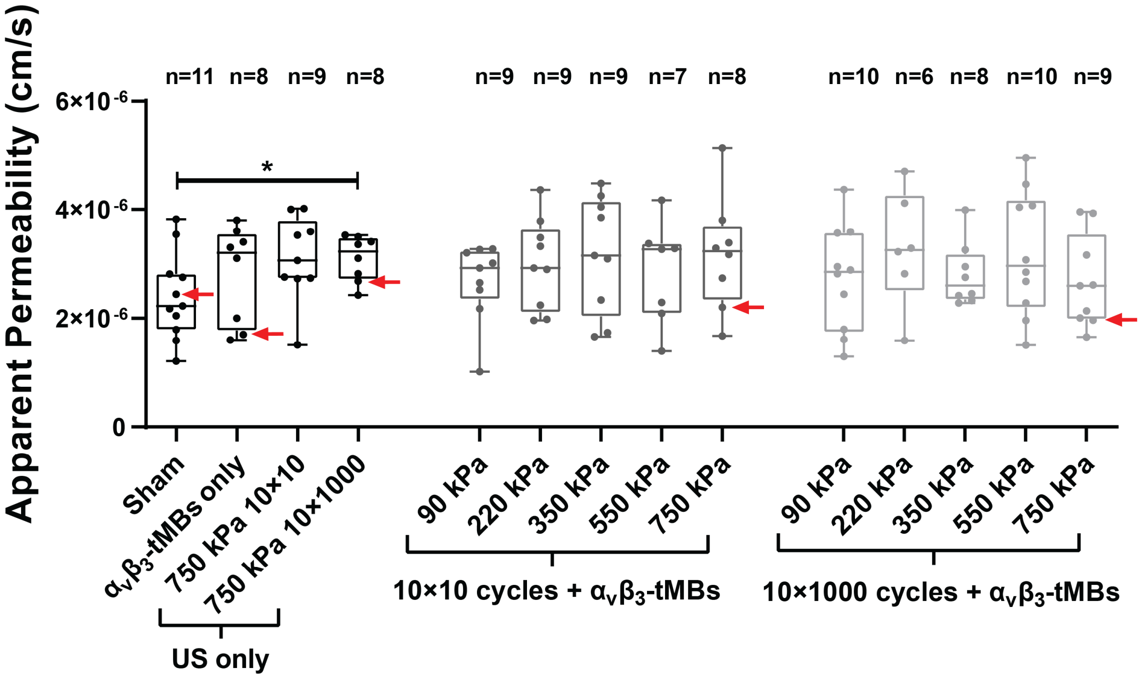


**Figure S4.** Apparent permeability before treatment in 121 microvessels for the 4 different control and 10 different ultrasound plus microbubble conditions. Boxplot represents the median and the boxes indicate the 25th and 75th percentiles with whiskers ranging from the minimum to maximum value. Statistical significance is indicated with *p<0.05, see Table S1 for all statistical comparisons. Datapoints which correspond to the example vessels shown in Figure 2 are indicated with red arrows. US = ultrasound; $\alpha_v\beta_3$-tMBs = $\alpha_v\beta_3$-targeted microbubbles.

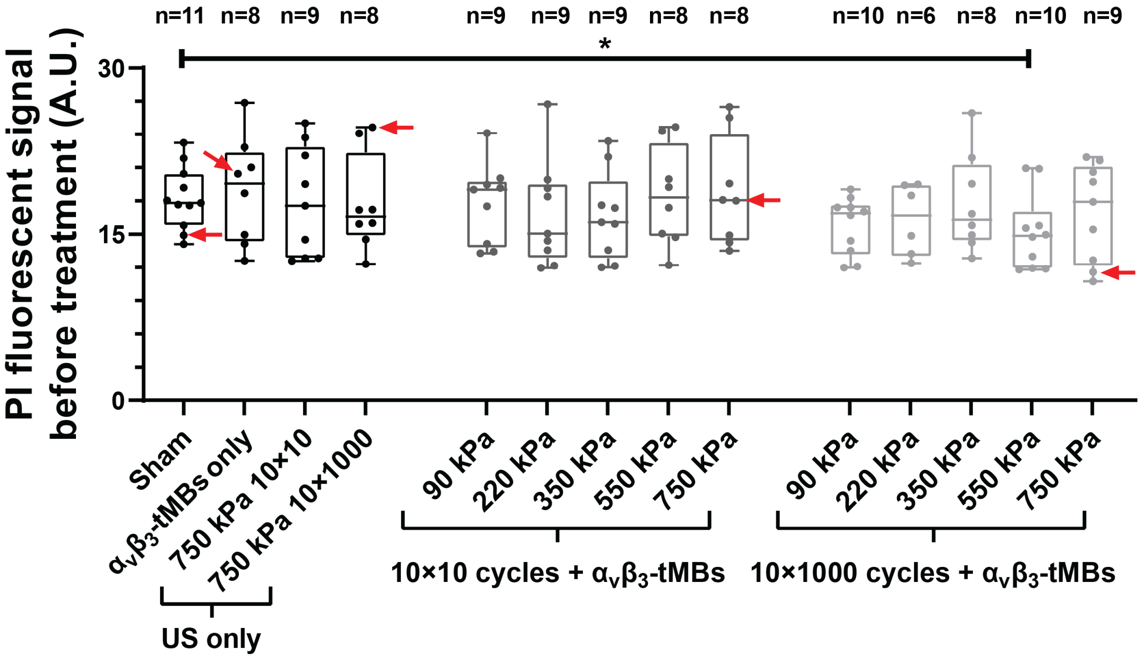


**Figure S5.** Quantification of the fluorescent propidium iodide signal before treatment. Boxplot represents the median and the boxes indicate the 25th and 75th percentiles with whiskers ranging from the minimum to maximum value. Statistical significance is indicated with *$p<0.05$, see Table S3 for all statistical comparisons. Datapoints which correspond to the example vessels shown in Figure 2 are indicated with red arrows. PI=propidium iodide; A.U. = arbitrary unit; US = ultrasound: $\alpha_v\beta_3$-tMBs = $\alpha_v\beta_3$-targeted microbubbles.

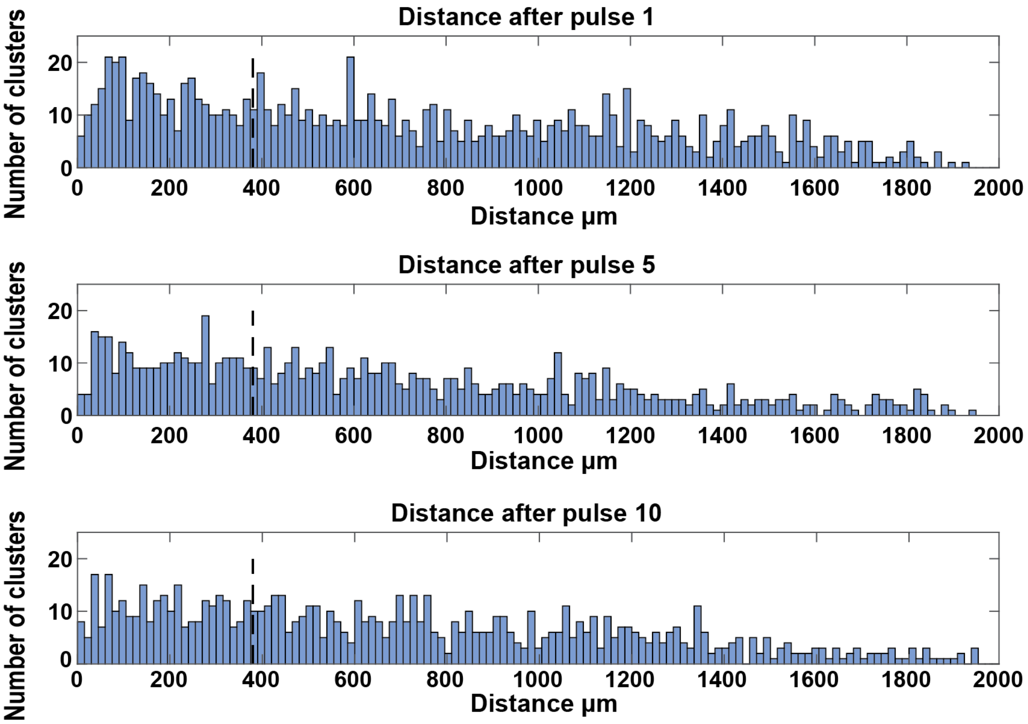


**Figure S6.** Quantification of the distance between $\alpha_v\beta_3$-tMB clusters after one, five, and ten cycles of ultrasound. The dashed vertical black line is the distance at which a node would occur if a standing wave would be present in the microvessel.

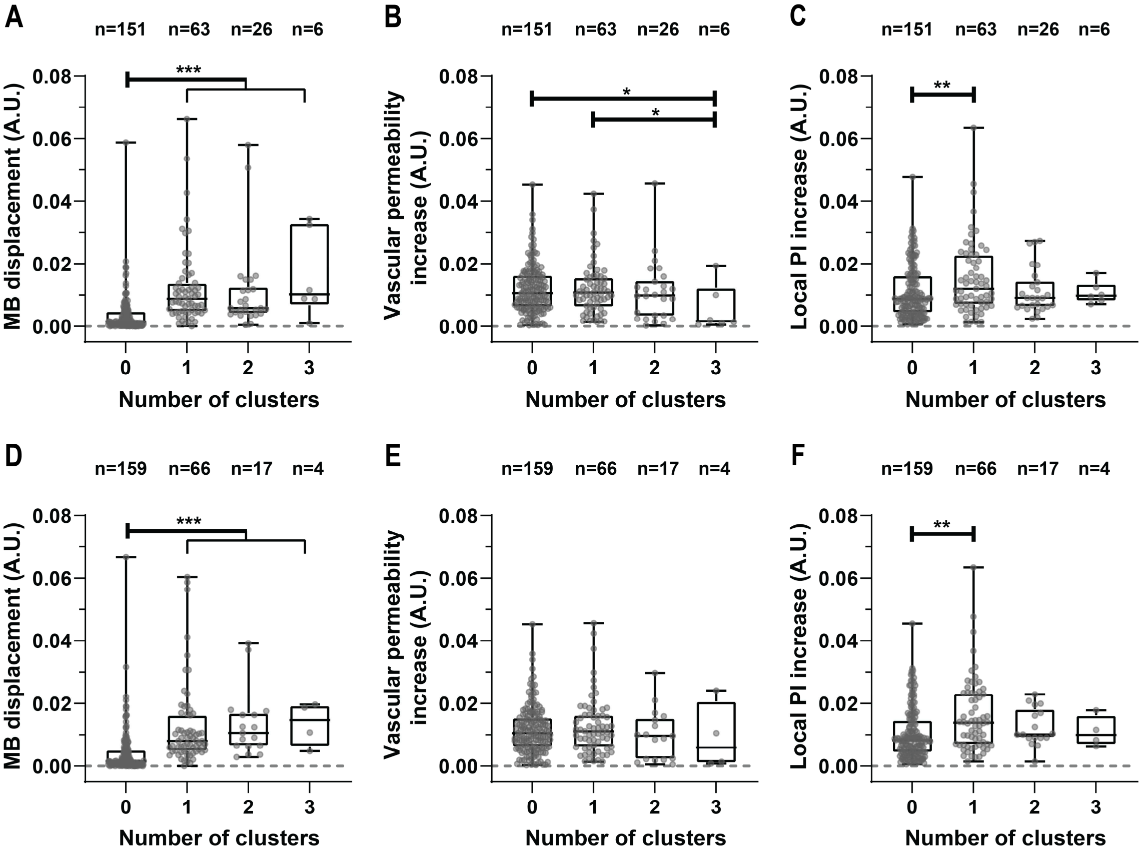


**Figure S7.** Local relations with the number of formed $\alpha_v\beta_3$-tMB clusters per 200 µm segments in the microvessels after the fifth and tenth cycle of ultrasound. A) Local relation between formed $\alpha_v\beta_3$-tMB clusters and $\alpha_v\beta_3$-tMB displacement after the fifth cycle of ultrasound. B) Local relation between formed $\alpha_v\beta_3$-tMB clusters and vascular permeability increase after the fifth cycle of ultrasound. C) Local relation between formed $\alpha_v\beta_3$-tMB clusters and propidium iodide (PI) increase (i.e., sonoporation) after the fifth cycle of ultrasound. D) Local relation between formed $\alpha_v\beta_3$-tMB clusters and $\alpha_v\beta_3$-tMB displacement after the tenth cycle of ultrasound. E) Local relation between formed $\alpha_v\beta_3$-tMB clusters and vascular permeability increase after the tenth cycle of ultrasound. F) Local relation between formed $\alpha_v\beta_3$-tMB clusters and propidium iodide (PI) increase (i.e., sonoporation) after the tenth cycle of ultrasound. (A-F) Boxplots represent the median and the boxes indicate the 25$^{th}$ to 75$^{th}$ percentiles with whiskers ranging from the minimum to maximum values and points representing individual segments. Statistical significance is indicated with *$p<0.05$, **$p<0.01$ and ***$p<0.001$. MB = $\alpha_v\beta_3$-targeted microbubble; A.U. = arbitrary unit; PI=propidium iodide.

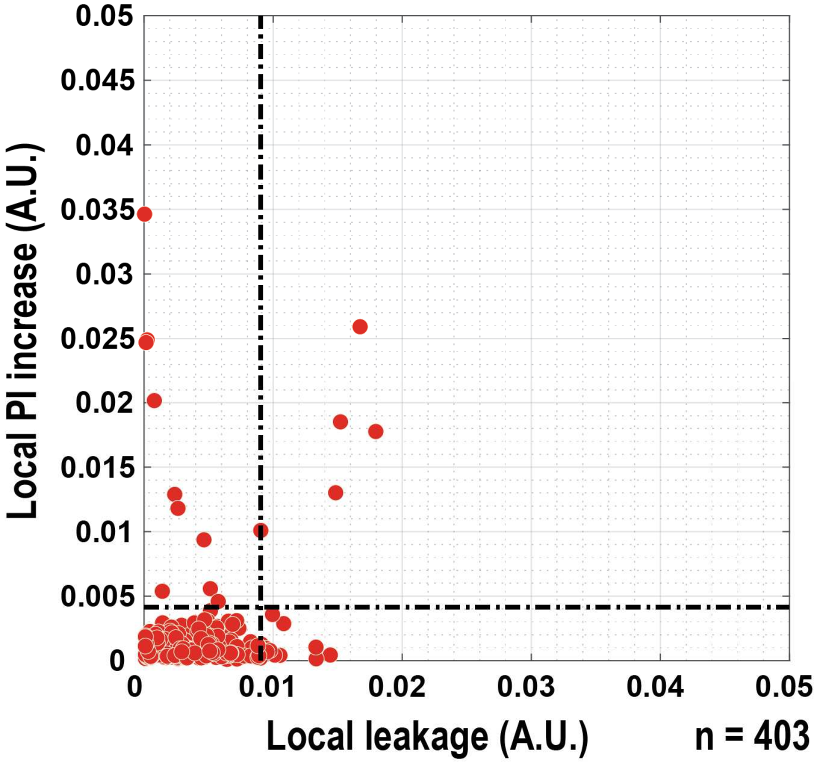


**Figure S8.** Correlation between vascular permeability increase (i.e., leakage) and sonoporation (i.e., propidium iodide uptake) in the microvessels for the four control treatments: sham, $\alpha_v\beta_3$-tMB only, 750 kPa 10×10 cycles ultrasound only, and 750 kPa 10×1000 cycles ultrasound only. Every data point and n-number represent individual microvessel segments of 200 µm. The dotted lines indicate the determined thresholds. PI=propidium iodide; A.U. = arbitrary unit.

| Apparent permeability before treatment | | Control groups | | | | 10 × 10 cycles + αvβ3-tMBs | | | | | 10 × 1000 cycles + αvβ3-tMBs | | | | |
|---|---|---|---|---|---|---|---|---|---|---|---|---|---|---|---|
| | | Sham | $\alpha_v\beta_3$-tMBs only | 750 kPa 10x10 US only | 750 kPa 10x1000 US only | 90 kPa | 220 kPa | 350 kPa | 550 kPa | 750 kPa | 90 kPa | 220 kPa | 350 kPa | 550 kPa | 750 kPa |
| Control groups | Sham | | 0.304 | 0.063 | 0.036 | 0.439 | 0.119 | 0.077 | 0.302 | 0.083 | 0.399 | 0.07 | 0.242 | 0.08 | 0.472 |
| | $\alpha_v\beta_3$-tMBs only | 0.304 | | 0.486 | 0.421 | 0.729 | 0.668 | 0.458 | 0.954 | 0.468 | 0.86 | 0.384 | 0.981 | 0.459 | 0.741 |
| | 750 kPa 10x10 US only | 0.063 | 0.486 | | 0.995 | 0.243 | 0.779 | 0.867 | 0.553 | 0.875 | 0.376 | 0.706 | 0.391 | 0.857 | 0.277 |
| | 750 kPa 10x1000 US only | 0.036 | 0.421 | 0.995 | | 0.16 | 0.748 | 0.851 | 0.49 | 0.856 | 0.331 | 0.659 | 0.259 | 0.844 | 0.211 |
| 10 × 10 cycles + αvβ3-tMBs | 90 kPa | 0.439 | 0.729 | 0.243 | 0.16 | | 0.391 | 0.255 | 0.692 | 0.261 | 0.88 | 0.204 | 0.689 | 0.263 | 0.996 |
| | 220 kPa | 0.119 | 0.668 | 0.779 | 0.748 | 0.391 | | 0.689 | 0.732 | 0.698 | 0.536 | 0.561 | 0.594 | 0.683 | 0.422 |
| | 350 kPa | 0.077 | 0.458 | 0.867 | 0.851 | 0.255 | 0.689 | | 0.52 | 0.994 | 0.353 | 0.849 | 0.392 | 0.987 | 0.279 |
| | 550 kPa | 0.302 | 0.954 | 0.553 | 0.49 | 0.692 | 0.732 | 0.52 | | 0.529 | 0.823 | 0.441 | 0.927 | 0.52 | 0.71 |
| | 750 kPa | 0.083 | 0.468 | 0.875 | 0.856 | 0.261 | 0.698 | 0.994 | 0.529 | | 0.366 | 0.846 | 0.394 | 0.982 | 0.288 |
| 10 × 1000 cycles + αvβ3-tMBs | 90 kPa | 0.399 | 0.86 | 0.376 | 0.331 | 0.88 | 0.536 | 0.353 | 0.823 | 0.366 | | 0.304 | 0.859 | 0.35 | 0.883 |
| | 220 kPa | 0.07 | 0.384 | 0.706 | 0.659 | 0.204 | 0.561 | 0.849 | 0.441 | 0.846 | 0.304 | | 0.298 | 0.864 | 0.234 |
| | 350 kPa | 0.242 | 0.981 | 0.391 | 0.259 | 0.689 | 0.594 | 0.392 | 0.927 | 0.394 | 0.859 | 0.298 | | 0.402 | 0.716 |
| | 550 kPa | 0.08 | 0.459 | 0.857 | 0.844 | 0.263 | 0.683 | 0.987 | 0.52 | 0.982 | 0.35 | 0.864 | 0.402 | | 0.283 |
| | 750 kPa | 0.472 | 0.741 | 0.277 | 0.211 | 0.996 | 0.422 | 0.279 | 0.71 | 0.288 | 0.883 | 0.234 | 0.716 | 0.283 | |

**Table S1.** Statistical comparison of the apparent permeability before treatment with the p-values per relation. The significant p=values of $p < 0.05$ (* in the Figures) are highlighted in green.

| Apparent permeability after treatment | | Control groups | | | | 10 × 10 cycles + αvβ3-tMBs | | | | | 10 × 1000 cycles + αvβ3-tMBs | | | | |
|---|---|---|---|---|---|---|---|---|---|---|---|---|---|---|---|
| | | Sham | $\alpha_v\beta_3$-tMBs only | 750 kPa 10x10 US only | 750 kPa 10x1000 US only | 90 kPa | 220 kPa | 350 kPa | 550 kPa | 750 kPa | 90 kPa | 220 kPa | 350 kPa | 550 kPa | 750 kPa |
| Control groups | Sham | | 1 | 0.412 | 0.002 | 0.603 | 0.941 | 0.503 | 0.117 | 0.0001 | 0.863 | 0.462 | 0.009 | 0.0000 | 0.0000 |
| | $\alpha_v\beta_3$-tMBs only | 1 | | 0.423 | 0.021 | 0.606 | 0.963 | 0.606 | 0.279 | 0.0010 | 0.965 | 0.414 | 0.028 | 0.0003 | 0.0001 |
| | 750 kPa 10x10 US only | 0.412 | 0.423 | | 0.321 | 0.666 | 0.297 | 0.605 | 0.888 | 0.0040 | 0.278 | 0.864 | 0.059 | 0.0010 | 0.0000 |
| | 750 kPa 10x1000 US only | 0.002 | 0.021 | 0.321 | | 0.008 | 0.027 | 0.027 | 0.015 | 0.0100 | 0.016 | 0.414 | 0.234 | 0.0040 | 0.0001 |
| 10 × 10 cycles + αvβ3-tMBs | 90 kPa | 0.603 | 0.606 | 0.666 | 0.008 | | 0.796 | 0.863 | 0.423 | 0.0003 | 0.4 | 0.864 | 0.021 | 0.0002 | 0.0000 |
| | 220 kPa | 0.941 | 0.963 | 0.297 | 0.027 | 0.796 | | 0.73 | 0.277 | 0.0006 | 0.842 | 0.388 | 0.015 | 0.0003 | 0.0000 |
| | 350 kPa | 0.503 | 0.606 | 0.605 | 0.027 | 0.863 | 0.73 | | 0.815 | 0.0010 | 0.604 | 0.864 | 0.021 | 0.0003 | 0.0000 |
| | 550 kPa | 0.117 | 0.279 | 0.888 | 0.015 | 0.423 | 0.277 | 0.815 | | 0.0010 | 0.146 | 0.852 | 0.065 | 0.0005 | 0.0001 |
| | 750 kPa | 0.0001 | 0.0010 | 0.0040 | 0.0100 | 0.0003 | 0.0006 | 0.0010 | 0.0010 | | 0.00032 | 0.008 | 0.721 | 0.2370 | 0.0460 |
| 10 × 1000 cycles + αvβ3-tMBs | 90 kPa | 0.863 | 0.965 | 0.278 | 0.016 | 0.4 | 0.842 | 0.604 | 0.146 | 0.00032 | | 0.428 | 0.012 | 0.0001 | 0.0000 |
| | 220 kPa | 0.462 | 0.414 | 0.864 | 0.414 | 0.864 | 0.388 | 0.864 | 0.852 | 0.008 | 0.428 | | 0.181 | 0.0070 | 0.0008 |
| | 350 kPa | 0.009 | 0.028 | 0.059 | 0.234 | 0.021 | 0.015 | 0.021 | 0.065 | 0.721 | 0.012 | 0.181 | | 0.1010 | 0.0150 |
| | 550 kPa | 0.0000 | 0.0003 | 0.0010 | 0.0040 | 0.0002 | 0.0003 | 0.0003 | 0.0005 | 0.2370 | 0.0001 | 0.0070 | 0.1010 | | 0.0950 |
| | 750 kPa | 0.0000 | 0.0001 | 0.0000 | 0.0001 | 0.0000 | 0.0000 | 0.0000 | 0.0001 | 0.0460 | 0.0000 | 0.0008 | 0.0150 | 0.0950 | |

**Table S2.** Statistical comparison of the apparent permeability after treatment with the p-values per relation. The significant p=values are highlighted in green: $p < 0.05$ (* in the Figures) is indicated with the lightest green color, $p < 0.01$ (** in the Figures) is indicated with the middle green color, and $p < 0.001$ (*** in the Figures) is indicated with the darkest green color.

| Fluorescent PI signal before treatment | | Control groups | | | | 10 × 10 cycles + αvβ3-tMBs | | | | | 10 × 1000 cycles + αvβ3-tMBs | | | | |
|---|---|---|---|---|---|---|---|---|---|---|---|---|---|---|---|
| | | Sham | $\alpha_v\beta_3$-tMBs only | 750 kPa 10x10 US only | 750 kPa 10x1000 US only | 90 kPa | 220 kPa | 350 kPa | 550 kPa | 750 kPa | 90 kPa | 220 kPa | 350 kPa | 550 kPa | 750 kPa |
| Control groups | Sham | | 0.778 | 0.71 | 0.492 | 0.766 | 0.331 | 0.261 | 1 | 0.968 | 0.051 | 0.404 | 0.6 | 0.036 | 0.71 |
| | $\alpha_v\beta_3$-tMBs only | 0.778 | | 0.673 | 0.721 | 0.606 | 0.277 | 0.37 | 0.878 | 0.798 | 0.122 | 0.228 | 0.721 | 0.122 | 0.481 |
| | 750 kPa 10x10 US only | 0.71 | 0.673 | | 1 | 0.931 | 0.666 | 0.546 | 0.815 | 0.481 | 0.315 | 0.607 | 0.963 | 0.278 | 0.605 |
| | 750 kPa 10x1000 US only | 0.492 | 0.721 | 1 | | 0.743 | 0.606 | 0.743 | 0.645 | 0.574 | 0.633 | 0.852 | 0.878 | 0.101 | 0.815 |
| 10 × 10 cycles + αvβ3-tMBs | 90 kPa | 0.766 | 0.606 | 0.931 | 0.743 | | 0.546 | 0.436 | 0.743 | 0.673 | 0.133 | 0.456 | 1 | 0.182 | 0.863 |
| | 220 kPa | 0.331 | 0.277 | 0.666 | 0.606 | 0.546 | | 1 | 0.37 | 0.606 | 0.604 | 0.955 | 0.606 | 0.497 | 0.863 |
| | 350 kPa | 0.261 | 0.37 | 0.546 | 0.743 | 0.436 | 1 | | 0.37 | 0.321 | 0.842 | 0.955 | 0.815 | 0.182 | 1 |
| | 550 kPa | 1 | 0.878 | 0.815 | 0.645 | 0.743 | 0.37 | 0.37 | | 0.959 | 0.173 | 0.491 | 0.798 | 0.146 | 0.673 |
| | 750 kPa | 0.968 | 0.798 | 0.481 | 0.574 | 0.673 | 0.606 | 0.321 | 0.959 | | 0.203 | 0.414 | 0.798 | 0.146 | 0.606 |
| 10 × 1000 cycles + αvβ3-tMBs | 90 kPa | 0.051 | 0.122 | 0.315 | 0.633 | 0.133 | 0.604 | 0.842 | 0.173 | 0.203 | | 0.492 | 0.515 | 0.393 | 0.497 |
| | 220 kPa | 0.404 | 0.228 | 0.607 | 0.852 | 0.456 | 0.955 | 0.955 | 0.491 | 0.414 | 0.492 | | 0.491 | 0.562 | 0.689 |
| | 350 kPa | 0.6 | 0.721 | 0.963 | 0.878 | 1 | 0.606 | 0.815 | 0.798 | 0.798 | 0.515 | 0.491 | | 0.173 | 0.815 |
| | 550 kPa | 0.036 | 0.122 | 0.278 | 0.101 | 0.182 | 0.497 | 0.182 | 0.146 | 0.146 | 0.393 | 0.562 | 0.173 | | 0.549 |
| | 750 kPa | 0.71 | 0.481 | 0.605 | 0.815 | 0.863 | 0.863 | 1 | 0.673 | 0.606 | 0.497 | 0.689 | 0.815 | 0.549 | |

**Table S3.** Statistical comparison of the fluorescent propidium iodide signal before treatment with the p-values per relation. The significant p=values of $p < 0.05$ (* in the Figures) are highlighted in green.

| Percentage change in PI signal within 45 s after treatment | | Control groups | | | | 10 × 10 cycles + αvβ3-tMBs | | | | | 10 × 1000 cycles + αvβ3-tMBs | | | | |
|---|---|---|---|---|---|---|---|---|---|---|---|---|---|---|---|
| | | Sham | $\alpha_v\beta_3$-tMBs only | 750 kPa 10x10 US only | 750 kPa 10x1000 US only | 90 kPa | 220 kPa | 350 kPa | 550 kPa | 750 kPa | 90 kPa | 220 kPa | 350 kPa | 550 kPa | 750 kPa |
| Control groups | Sham | | 0.84 | 0.503 | 0.442 | 0.331 | 0.175 | 0.046 | 0.026 | 0.009 | 0.282 | 0.007 | 0.003 | 0.0000 | 0.0000 |
| | $\alpha_v\beta_3$-tMBs only | 0.84 | | 0.481 | 0.234 | 0.2 | 0.093 | 0.046 | 0.028 | 0.015 | 0.237 | 0.02 | 0.005 | 0.0000 | 0.0001 |
| | 750 kPa 10x10 US only | 0.503 | 0.481 | | 0.888 | 0.73 | 0.489 | 0.222 | 0.114 | 0.027 | 1 | 0.026 | 0.015 | 0.0000 | 0.0000 |
| | 750 kPa 10x1000 US only | 0.442 | 0.234 | 0.888 | | 0.888 | 0.481 | 0.236 | 0.083 | 0.028 | 0.762 | 0.043 | 0.01 | 0.0000 | 0.0001 |
| 10 × 10 cycles + αvβ3-tMBs | 90 kPa | 0.331 | 0.2 | 0.73 | 0.888 | | 0.489 | 0.222 | 0.074 | 0.021 | 0.842 | 0.05 | 0.011 | 0.0000 | 0.0000 |
| | 220 kPa | 0.175 | 0.093 | 0.489 | 0.481 | 0.489 | | 0.546 | 0.277 | 0.036 | 0.72 | 0.05 | 0.021 | 0.0000 | 0.0000 |
| | 350 kPa | 0.046 | 0.046 | 0.222 | 0.236 | 0.222 | 0.546 | | 0.423 | 0.059 | 0.4 | 0.113 | 0.027 | 0.0000 | 0.0000 |
| | 550 kPa | 0.026 | 0.028 | 0.114 | 0.083 | 0.074 | 0.277 | 0.423 | | 0.382 | 0.203 | 0.491 | 0.05 | 0.0001 | 0.0001 |
| | 750 kPa | 0.009 | 0.015 | 0.027 | 0.028 | 0.021 | 0.036 | 0.059 | 0.382 | | 0.027 | 0.95 | 0.161 | 0.0003 | 0.0001 |
| 10 × 1000 cycles + αvβ3-tMBs | 90 kPa | 0.282 | 0.237 | 1 | 0.762 | 0.842 | 0.72 | 0.4 | 0.203 | 0.027 | | 0.031 | 0.021 | 0.0000 | 0.0000 |
| | 220 kPa | 0.007 | 0.02 | 0.026 | 0.043 | 0.05 | 0.05 | 0.113 | 0.491 | 0.95 | 0.031 | | 0.081 | 0.0010 | 0.0004 |
| | 350 kPa | 0.003 | 0.005 | 0.015 | 0.01 | 0.011 | 0.021 | 0.027 | 0.05 | 0.161 | 0.021 | 0.081 | | 0.0060 | 0.0010 |
| | 550 kPa | 0.0000 | 0.0000 | 0.0000 | 0.0000 | 0.0000 | 0.0000 | 0.0000 | 0.0001 | 0.0003 | 0.0000 | 0.0010 | 0.0060 | | 0.6610 |
| | 750 kPa | 0.0000 | 0.0001 | 0.0000 | 0.0001 | 0.0000 | 0.0000 | 0.0000 | 0.0001 | 0.0001 | 0.0000 | 0.0004 | 0.0010 | 0.6610 | |

**Table S4.** Statistical comparison of change in fluorescent propidium iodide signal after treatment with the p-values per relation. The significant p=values are highlighted in green: $p < 0.05$ (* in the Figures) is indicated with the lightest green color, $p < 0.01$ (** in the Figures) is indicated with the middle green color, and $p < 0.001$ (*** in the Figures) is indicated with the darkest green color.

| | Number of clusters per condition | | | | | |
|---|---|---|---|---|---|---|
| **Pressure (kPa)** / **Cycles** | 90 | 220 | 350 | 550 | 750 | Total |
| 10x10 | 0 | 1 | 3 | 0 | 0 | 4 |
| 10x1000 | 4 | 8 | 16 | 92 | 87 | 207 |

**Table S5.** Number of clusters per condition after the first cycle of ultrasound.

| WST-8 cell viability assay after treatment | | WST-8 assay control groups | | | | | Control groups | | | | 10 × 10 cycles + | | 10 × 1000 cycles + αvβ3- | | |
|---|---|---|---|---|---|---|---|---|---|---|---|---|---|---|---|
| | | Cell free | Without WST-8 reagent | Positive control (dead cells) | Cell free + αvβ3-tMBs | Cell free + 850 kPa 10x1000 + αvβ3-tMBs | Sham | $\alpha_v\beta_3$-tMBs only | 750 kPa 10x10 US only | 750 kPa 10x1000 US only | 550 kPa | 750 kPa | 350 kPa | 550 kPa | 750 kPa |
| WST-8 assay control groups | Cell free | | 0.61 | 0.31 | 0.714 | 0.905 | 0.00067 | 0.002 | 0.001 | 0.000666 | 0.002 | 0.002 | 0.002 | 0.002 | 0.001 |
| | Without WST-8 reagent | 0.61 | | 0.476 | 0.629 | 0.629 | 0.004 | 0.01 | 0.006 | 0.004 | 0.01 | 0.01 | 0.01 | 0.01 | 0.006 |
| | Positive control (dead cells) | 0.31 | 0.476 | | 0.548 | 0.381 | 0.00067 | 0.002 | 0.001 | 0.000666 | 0.002 | 0.002 | 0.002 | 0.002 | 0.001 |
| | Cell free + αvβ3-tMBs | 0.714 | 0.629 | 0.548 | | 1 | 0.012 | 0.024 | 0.017 | 0.012 | 0.024 | 0.024 | 0.024 | 0.024 | 0.017 |
| | Cell free + 850 kPa 10x1000 + αvβ3-tMBs | 0.905 | 0.629 | 0.381 | 1 | | 0.012 | 0.024 | 0.017 | 0.012 | 0.024 | 0.024 | 0.024 | 0.024 | 0.017 |
| Control groups | Sham | 0.00067 | 0.004 | 0.00067 | 0.012 | 0.012 | | 0.414 | 0.694 | 0.505 | 0.852 | 0.662 | 0.414 | 0.345 | 0.397 |
| | $\alpha_v\beta_3$-tMBs only | 0.002 | 0.01 | 0.002 | 0.024 | 0.024 | 0.414 | | 0.234 | 0.755 | 0.394 | 0.818 | 0.937 | 1 | 0.836 |
| | 750 kPa 10x10 US only | 0.001 | 0.006 | 0.001 | 0.017 | 0.017 | 0.694 | 0.234 | | 0.397 | 0.534 | 0.234 | 0.234 | 0.101 | 0.128 |
| | 750 kPa 10x1000 US only | 0.00067 | 0.004 | 0.00067 | 0.012 | 0.012 | 0.505 | 0.755 | 0.397 | | 0.414 | 0.755 | 1 | 0.95 | 0.955 |
| 10 × 10 cycles + αvβ3-tMBs | 550 kPa | 0.002 | 0.01 | 0.002 | 0.024 | 0.024 | 0.852 | 0.394 | 0.534 | 0.414 | | 0.818 | 0.394 | 0.18 | 0.534 |
| | 750 kPa | 0.002 | 0.01 | 0.002 | 0.024 | 0.024 | 0.662 | 0.818 | 0.234 | 0.755 | 0.818 | | 0.818 | 0.394 | 0.836 |
| 10 × 1000 cycles + αvβ3-tMBs | 350 kPa | 0.002 | 0.01 | 0.002 | 0.024 | 0.024 | 0.414 | 0.937 | 0.234 | 1 | 0.394 | 0.818 | | 1 | 0.836 |
| | 550 kPa | 0.002 | 0.01 | 0.002 | 0.024 | 0.024 | 0.345 | 1 | 0.101 | 0.95 | 0.18 | 0.394 | 1 | | 0.445 |
| | 750 kPa | 0.001 | 0.006 | 0.001 | 0.017 | 0.017 | 0.397 | 0.836 | 0.128 | 0.955 | 0.534 | 0.836 | 0.836 | 0.445 | |

**Table S6.** Statistical comparison of the WST-8 cell viability assay after treatment with the p-values per relation. The significant p=values are highlighted in green: $p < 0.05$ (* in the Figures) is indicated with the lightest green color, $p < 0.01$ (** in the Figures) is indicated with the middle green color, and $p < 0.001$ (*** in the Figures) is indicated with the darkest green color.
